# Characterising a species-rich and understudied tropical insect fauna using DNA barcoding

**DOI:** 10.1101/2025.09.30.675770

**Authors:** David R. Hemprich-Bennett, Ezekiel Donkor, Bernard Adams, Naana Afua Acquaah, Eva D. Ofori, Samuel Anie-Amoah, Abigail Bailey, H. Charles J. Godfray, Owen T. Lewis, Fred Aboagye-Antwi, Talya D. Hackett

## Abstract

**Background:** West Africa has high biodiversity that is relatively understudied, especially for insects. Studies of West African arthropod diversity can therefore help address important questions regarding conservation, ecosystem services, and insecticide use and other species-control interventions in agriculture and disease management. We intensively sampled arthropods in Ghana using complementary trapping methods, generated DNA barcodes, and classified sequences by Barcode Index Numbers (BINs, a species proxy). Using this dataset, we investigate assemblage composition, temporal activity patterns, and the state of regional biodiversity sampling.

**Results:** Sequencing DNA from 95,996 individuals captured using Malaise, yellow pan, pitfall, Heath and Centre for Disease Control (CDC) traps, we identified 10,120 unique BINs. The rate of species accumulation did not approach an asymptote for any taxonomic group or trap type, indicating high biodiversity. The different trap types sampled different subsets of the local community, with greatest similarity between yellow pan and pitfall traps. More insects and species (BINs) were trapped during the day than at night. Our dataset shared more BINs in the Barcode of Life Database with South Africa than with any other country, although this predominantly reflects the limited sampling and DNA sequencing campaigns in Africa.

**Conclusions:** This study more than doubles the published BINs for West Africa, offering insights into the biodiversity of an ecologically important but understudied taxon and region. Using multiple trap types allowed a more complete assessment of the local arthropod assemblage. The public release of these data will support and stimulate further taxonomic and ecological work in the region.

## Background

Tropical regions harbour approximately 50% of the world’s described species [1], but only 1 million out of an estimated 7 million species of terrestrial arthropods (predominantly insects) have been formally described [2]. The global dry biomass of all insects is approximately 300 million metric tonnes [3], similar to the combined biomass of humanity and its livestock. Given their functional diversity and numerical dominance in many terrestrial ecosystems, arthropods play critical roles in global ecosystem functioning.

Documenting the extraordinary diversity of insect taxa is increasingly important because of the threats that tropical ecosystems are experiencing, including climate change, land-use change, introduced species and pollution [4]. These threats are expected to affect many species supra- additively, and with uneven impacts even for closely related taxa [5]. The impacts of these changes are better understood for more heavily studied taxa such as vertebrates [6]. Low levels of baseline data and taxonomic challenges limit our ability to measure changes in insect populations and communities. As a result, the few studies on insect declines tend to come from better-studied temperate faunas in Europe and North America [7], or from a few tropical sites where well-established field stations have facilitated relatively intensive and long-term study, e.g. [8]. A major impediment to tropical insect studies is that even a brief sampling campaign generates a huge number and diversity of specimens with concomitant logistical and taxonomic challenges [9].

These taxonomic and sampling impediments prevent our understanding of the impact of specific interventions, such as crop pest and disease vector control, on non-target insect populations and broader ecological communities. In particular, technological advances are generating new methods for species-specific targeting of pests and vectors thereby avoiding widespread biocide use [10–12]. Regulators and policy makers require an understanding of the wider biodiversity impacts of these new technologies.

In the past two decades, DNA barcoding has emerged as a key technique to accelerate rates of sample characterisation. DNA barcoding works by comparing specific regions of DNA from collected specimens to sequences in a reference database, enabling researchers to infer their likely identity [13]. A partial sequence of the cytochrome c oxidase subunit I (COI) gene has emerged as the near-universal barcode region for arthropods, being highly conserved within species, but variable between species. The primary repository for insect DNA barcodes is the Barcode of Life Data System (BOLD) [14] which stores publicly accessible sequences and associated metadata, and assigns a unique identifier to each sequence cluster, known as a Barcode Index Number (BIN) [15]. BINs are clusters of highly similar DNA sequences and so can act as a species-proxy when a species has yet to be formally described. BINs can thus be used in ecological studies where species names are not essential, for example in identifying biodiversity hotspots, estimating trends in abundance and diversity, or understanding ecological community structure. DNA barcodes are also an invaluable tool for species discovery, especially in areas where coverage of described species is high [16]. Uploading samples to BOLD provides a useful resource for future researchers, allowing samples to be matched to previous records along with associated taxonomic, geographic and temporal metadata. In addition, CO1 barcodes can be used in metabarcoding studies, acting as reference sequences for eDNA [17] or dietary analysis [18]. BOLD currently has 16.5 million public records, representing 1.2 million BINs.

Here we use BINs as Operational Taxonomic Units (OTUs) where samples are classified into clusters based on shared or diverging sequences. In this form, BINs have been used to analyse ecological patterns in the tropics [19–22], while simultaneously revealing the presence of many previously unknown taxa.

In this paper we describe the implementation and outcomes of a high-intensity arthropod- sampling campaign in a biodiversity-rich area in Ghana, West Africa. Like many tropical regions, West African forests and savannas have high levels of biodiversity that is threatened by human actions [23]. Compared even to other tropical regions, relatively few arthropod species in West Africa have been described, and species from this region are under-represented in global DNA datasets such as BOLD, hampering ecological study, estimates of species distribution, extinction and more [24].

Our insect sampling and DNA barcoding campaign primarily took place to make a reference library for use in an ongoing project investigating the position the mosquito malarial vector, *Anopheles gambiae* [25], in its local ecological community and to assess the effects of different control strategies on non-target organisms. The resulting dataset – which is publicly available and analysed here – will also provide a rich resource for biologists concerned with documenting arthropod diversity in West Africa.

Specifically, we ask: (1) What fraction of the local arthropod assemblage does our sample of nearly 100K individuals reveal? (2) How important is it to use multiple trap types to sample biodiversity? (3) How do insect activity patterns differ between day and night? (4) Which countries represented in the BOLD database share the most BINs with our dataset, and what does this tell us about the global completeness of the BOLD dataset? More broadly, the resulting data on rates of taxonomic discovery through time and across trapping methods will support optimising survey methods and strategies for other poorly studied but species-rich arthropod communities.

## Methods

### Field sites

Samples were collected at and adjacent to two villages (‘sites’) in the Volta region of Ghana, Abutia Amegame in the Ho West District (6.209 N, 0.441 E) and Mafi Agove in the Central Tongu District (6.457 N, 0.316 E). Monthly sampling campaigns were conducted from February 2019 to March 2020 inclusive, and (following a pause imposed by the COVID-19 pandemic) from April to June 2021. The villages are small subsistence farming settlements (<1,000 people) within a matrix of grassland, cropland and forest fragments, in a region characterised by pronounced annual dry and rainy seasons.

### Sampling

We sampled within a 500 m radius of village centres, dividing each circle into four equal quadrants (NW, NE, SE, SW). At each village, on every visit, we set four transects (one within each quadrant) at random, pre-determined start locations and directions (Supplementary Figure 1, Supplementary Figure 2). We placed a Malaise trap at the start of a transect orientated in a direction most likely to intersect with arthropod flight paths based on the typical wind direction, topography and vegetation. At 10 m, 20 m, 30 m, and 40 m, we placed a yellow pan trap and a pitfall trap, both filled with soapy water on alternating sides of the transect line. We set a Heath trap at 50 m and a CO_2_-baited Center for Disease Control (CDC) trap at the 100 m point to avoid interference with other traps. Traps were left for 24 hours, and Malaise trap bottles were exchanged at 06:00, 12:00, 18:00 and 00:00 to capture temporal dynamics during the sampling period. The arthropods collected using each trap type on each transect on each date is referred to as a ‘Lot’.

### DNA barcoding

We sub-sorted all Lots before they were sent for sequencing to maximise diversity. Individuals from Malaise, pan, pitfall and CDC Lots were identified to taxonomic order, assigned a morphospecies identity based on visual inspection, and up to five individuals per morphospecies per Lot were selected for sequencing. Heath trap Lots had considerably higher arthropod abundance and richness than those from other trap types. The number of arthropods selected for sequencing per Lot was in proportion to their wet mass, while aiming to capture the diversity of the Lot. Due to logistical and financial constraints only 34 out of 117 Malaise lots were fully sequenced. Araneae (spider) samples from pan and pitfall traps were removed for use in ongoing dietary analyses, and so Araneae are omitted from some analyses here.

Samples were DNA barcoded at the Canadian Centre for DNA Barcoding (CCDB), using their standard protocol: photographing each specimen before performing a non-destructive DNA extraction, PCR using the CO1 ‘Folmer’ region [26], and sequencing on a PacBio Sequel.

Through the BOLD data-management platform [14], samples were assigned provisional taxonomy (typically to family level, though more precise information was automatically assigned where possible) and a BIN [15]. Where species-level assignments were made, the resulting species list was queried against a custom database of pests of crops and human health, assembled from a manual search of the literature.

## Data analyses

For final analyses our data were downloaded from BOLD on 15th September 2025, and BOLD was at the same time queried for information on the publicly available BINs that matched BINs in our dataset. All analyses used R 4.5.1 [27], with plots created using the ggplot2 package [28]. All code used in this manuscript is available at https://github.com/hemprichbennett/ghana_bins.

### Species richness and sampling completeness

Sampling completeness was calculated for each taxonomic order and trap type using the ‘iNEXT’ R package [29] . The rate of BIN-accumulation was used to estimate how many individual insects it would be necessary to sequence to document all BINs that would be captured by a given trap-type. Calculations were restricted to combinations of taxonomic order and trap-type where at least 20 BINs were detected.

### Assemblage composition comparisons

To compare assemblages among trap types, a series of non-metric multidimensional scaling (NMDS) analyses were run using the ‘vegan’ R package [30] at order, family, genus and BIN levels. Taxonomic groups were only included in the analyses if they contained a minimum of 10 samples. Each sampling location was categorised by both trap type and habitat type (forest, semi-natural, or village). CDC traps were not included for the genus-level analyses, as there were insufficient samples assigned to the level of genus.

To test for differences among the assemblages included in the NMDS analyses, we also ran a Permutational Multivariate Analysis of Variance using Bray-Curtis dissimilarity values and tested for the interaction between trap type and habitat.

### Temporal analyses

To explore temporal changes in arthropod assemblages for the Malaise trap data we ran two linear mixed effects models. We modelled the change in number of insects captured (abundance) or the number of BINs detected (richness) with the time of day as a fixed effect and allowing a random intercept for the lot.

To compare diurnal and nocturnal assemblages we ran a set of NMDS and Permutational Multivariate Analysis of Variance analyses at order, family, genus and BIN levels, as above.

We tested if numbers of arthropod individuals and BINs captured were more variable in the daytime than at night with Brown-Forsythe tests using the ‘onewaytests’ package [31].

### Geographic analyses

We queried BOLD for the available metadata corresponding to all BINs in our dataset that already had public matches from other studies. The resulting dataset was used to identify countries and continents which shared BINs with our dataset, and the taxonomic assignment of those BINs. Data for the 20 countries with the highest numbers of public BINs matching BINs in our dataset were queried on September 15th, 2025, to investigate the extent to which similarity in arthropod assemblages is a result of geographic variations in sampling and sequencing effort versus proximity to Ghana. This linear regression analysis modelled the number of BINs shared with our dataset for each country as a function of the country’s geographic distance from Ghana and the number of public sequences for each country as fixed effects.

We expected the number of shared BINs to be more impacted by the country’s sampling effort than their distance from Ghana, so that, for example, the relatively well-sampled Costa Rica might have more BINs in common with our dataset than the under-sampled but neighbouring Togo.

## Results

### Overview of the barcode library

We recorded 10,120 unique BINs across the 95,996 samples sequenced (Table 1), of which 4,939 were newly recorded in our project. At the time of writing, this total represents ∼0.8% of the total BINs for all taxa on BOLD. Notably, only 8.5% of public BINs on BOLD originate from Africa, meaning our contribution accounts for nearly 10% of all African taxa in the database. Excluding Ghana, only 4,418 public BINs are from West Africa, and our dataset alone contributes more than double that number. Heath traps, with the greatest number of sequenced samples, provided the most novel BINs, (i.e., those not previously represented on BOLD) (2,850) (Table 1, Supplementary Figure 3).

**Table 1:** The number of samples sequenced and the number and diversity of BINs documented for each trap type

| **Trap type** | **Number of samples** | **Number of BINs** | **Number of BINs unique to the trap type** | **Shannon Diversity** |
| --- | --- | --- | --- | --- |
| CDC | 3,039 | 758 | 248 | 4.20 |
| Heath | 65,293 | 6,896 | 5,287 | 6.67 |
| Malaise | 11,975 | 3,233 | 1,810 | 6.40 |
| Pitfall | 8,296 | 856 | 232 | 4.20 |
| Yellow Pan | 7,223 | 1,345 | 444 | 4.90 |

We recorded 31 taxonomic orders and 384 taxonomic families, with 92% of samples being Coleoptera, Diptera, Hemiptera, Hymenoptera or Lepidoptera (Table 2, Supplementary Table 1, Supplementary Table 2). 583 samples (25 BINs) were known crop pest species Supplementary Table 3), with 264 samples belonging to the families of Diptera which are known to blood-feed (Culicidae, Simuliidae, Tabanidae); 192 were Culicidae, a hematophagous taxa of particular interest (Supplementary Table 4).

**Table 2:** The percentage constitution of each taxonomic order of samples sequenced per trap type. E.g. 57.85% of all samples sequenced from CDC traps were Dipterans.

| Order | Trap type |  |  |  |  |
| --- | --- | --- | --- | --- | --- |
|  | CDC (%) | Heath (%) | Malaise | Pitfall (%) | Yellow Pan (%) |
| Araneae | 0.49 | 0.21 | 0 | 0 | 0.03 |
| Archaeognatha | 0 | 0 | 0 | 0.05 | 0.06 |
| Blattodea | 0.07 | 0.6 | 0.33 | 1.25 | 0.8 |
| Coleoptera | 13.43 | 33.02 | 5.6 | 13.39 | 8.49 |
| Dermaptera | 0.07 | 0.11 | 0.04 | 0.13 | 0.08 |
| Diptera | 57.85 | 9.4 | 54.7 | 7.22 | 28.77 |
| Embioptera | 0 | 0.01 | 0.01 | 0 | 0 |
| Entomobryomorpha | 0.07 | 0.09 | 0.3 | 2.74 | 2.44 |
| Ephemeroptera | 0 | 0.33 | 0 | 0 | 0 |
| Hemiptera | 4.61 | 19.58 | 11.38 | 5.61 | 14.97 |
| Hymenoptera | 10.04 | 9.52 | 16.64 | 58.47 | 36.74 |
| Isopoda | 0 | 0 | 0 | 0.02 | 0 |
| Ixodida | 0 | 0 | 0 | 0.04 | 0 |
| Lepidoptera | 11.19 | 22.62 | 6.82 | 0.22 | 0.57 |
| Mantodea | 0.03 | 0.06 | 0 | 0.04 | 0.06 |
| Mecoptera | 0 | 0.02 | 0 | 0 | 0 |
| Mesostigmata | 0 | 0.09 | 0 | 0 | 0 |
| Neuroptera | 0 | 0.17 | 0.16 | 0.05 | 0.03 |
| Odonata | 0.07 | 0.01 | 0.03 | 0 | 0.03 |
| Orthoptera | 0.43 | 2.7 | 1.84 | 9.99 | 5.39 |
| Phasmida | 0 | 0 | 0 | 0.01 | 0 |
| Plecoptera | 0.1 | 0 | 0.01 | 0.01 | 0.01 |
| Poduromorpha | 0 | 0 | 0 | 0.01 | 0 |
| Pseudoscorpiones | 0 | 0.01 | 0 | 0 | 0 |
| Psocodea | 0.59 | 0.13 | 0.82 | 0.1 | 0.26 |
| Strepsiptera | 0 | 0.03 | 0 | 0 | 0 |
| Symphyleona | 0 | 0.01 | 0.03 | 0 | 0.04 |
| Thysanoptera | 0.07 | 0.21 | 0.03 | 0.1 | 0.75 |
| Trichoptera | 0 | 0.7 | 0.94 | 0.01 | 0.04 |

### BIN accumulation

For the 13 most abundant taxonomic orders, the average sampling completeness (the percentage of BINs detected relative to those estimated to exist within the community) was 53.3%.

Neuroptera had the lowest completeness at 13.5%, while Trichoptera had the highest at 71.1%. None of the trap types or taxonomic orders reached full completeness (Figure 1), and some orders such as Coleoptera, Diptera, and Lepidoptera were estimated to have thousands of unsampled BINs present at our sites.

**Figure 1:**
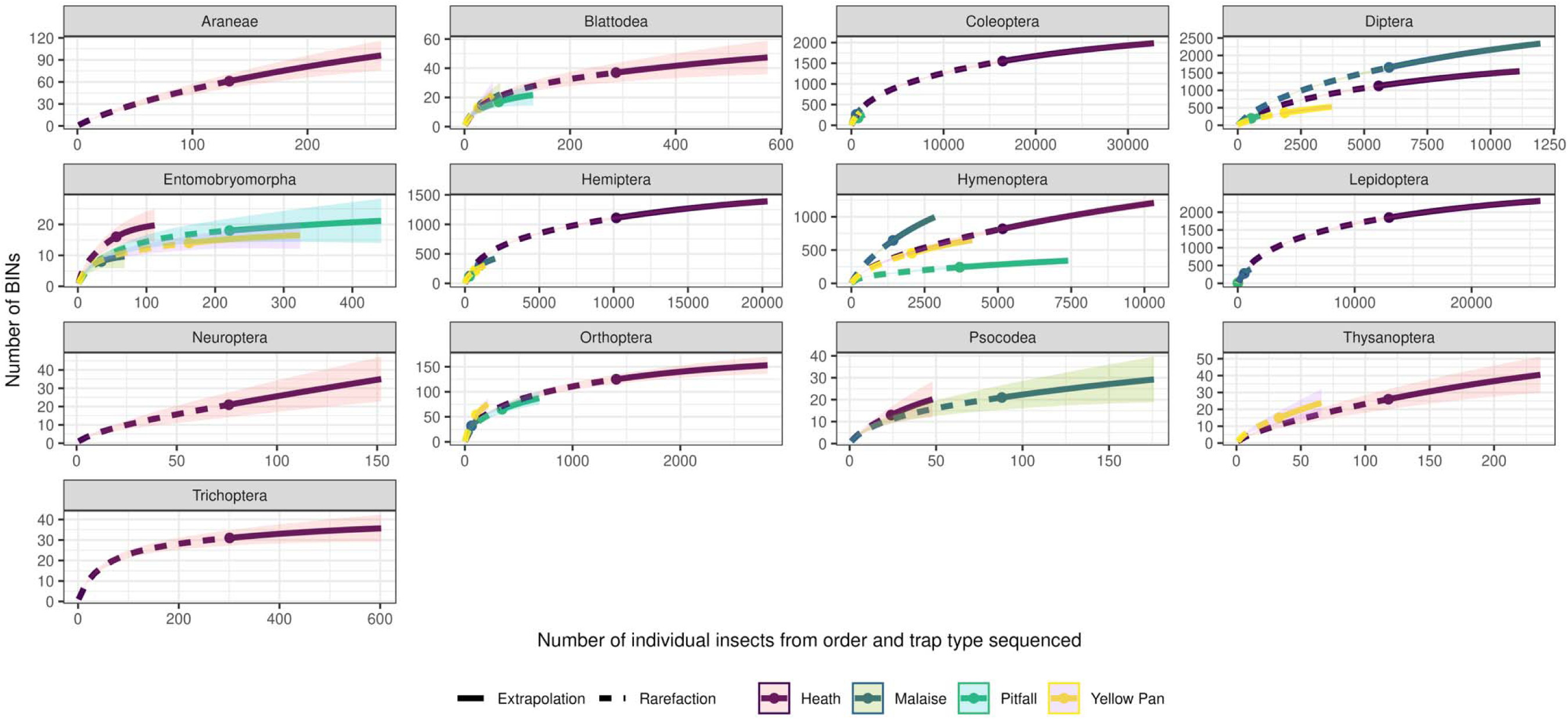
Type 1 iNEXT plots for different taxa showing the observed and extrapolated accumulation of BINs in relation to the number of individuals sequenced. Curves are plotted separately for each trapping method.

### Trap complementarity

Overall, assemblages of arthropods caught in each trap type differed significantly at the level of order (F_4,2_ = 171.8, p<0.001, R^2^ = 0.46), family (F_4,2_ = 106.3, p < 0.001, R^2^ = 0.36), genus (F_3,2_=6.0, p<0.001, R^2^ = 0.17) and BIN (F_4,2_ = 22.9, p<0.001, R^2^ = 0.12). The degree of overlap among trap types decreased with increasing taxonomic resolution (Figure 2: NMDS plots comparing insect assemblages among trap types. Each plot differs in terms of the taxonomic resolution at which samples were categorised. The larger points are the centroids for each trap type.Figure 2), showing the advantage of fine-scale taxonomic IDs (in this case BINs) when analysing diverse arthropod assemblages. Heath traps were distinct from the other trap types, even at the order level, but there was near-complete order-level overlap between pan and pitfall traps (Figure 2a). However, at the BIN-level, there was very little overlap, with each trap type capturing a distinct arthropod assemblage (Figure 2d), highlighting the importance of using multiple trap types when surveying arthropods to gain a more representative dataset. Due to the great richness of taxa captured there were few clear trends of specific taxonomic groups particularly driving these differences, but in general the Culicidae were primarily captured in CDC traps, Formicidae in Heath and pitfall traps, Chlopidae and Muscidae in Malaise traps, and Dolichopodidae mostly in yellow pan traps. Reflecting their relative distinctness from the other trap types, several taxa were predominantly captured in Heath traps, including Termitidae, Carabidae, Chrysomelidae, Dysticidae, Scarabidae, Staphylinidae, Cicadellidae, Miridae, Rhyparochromidae, Braconidae, Lepidoptera, Mantodea, Orthoptera and Trichoptera.

**Figure 2:**
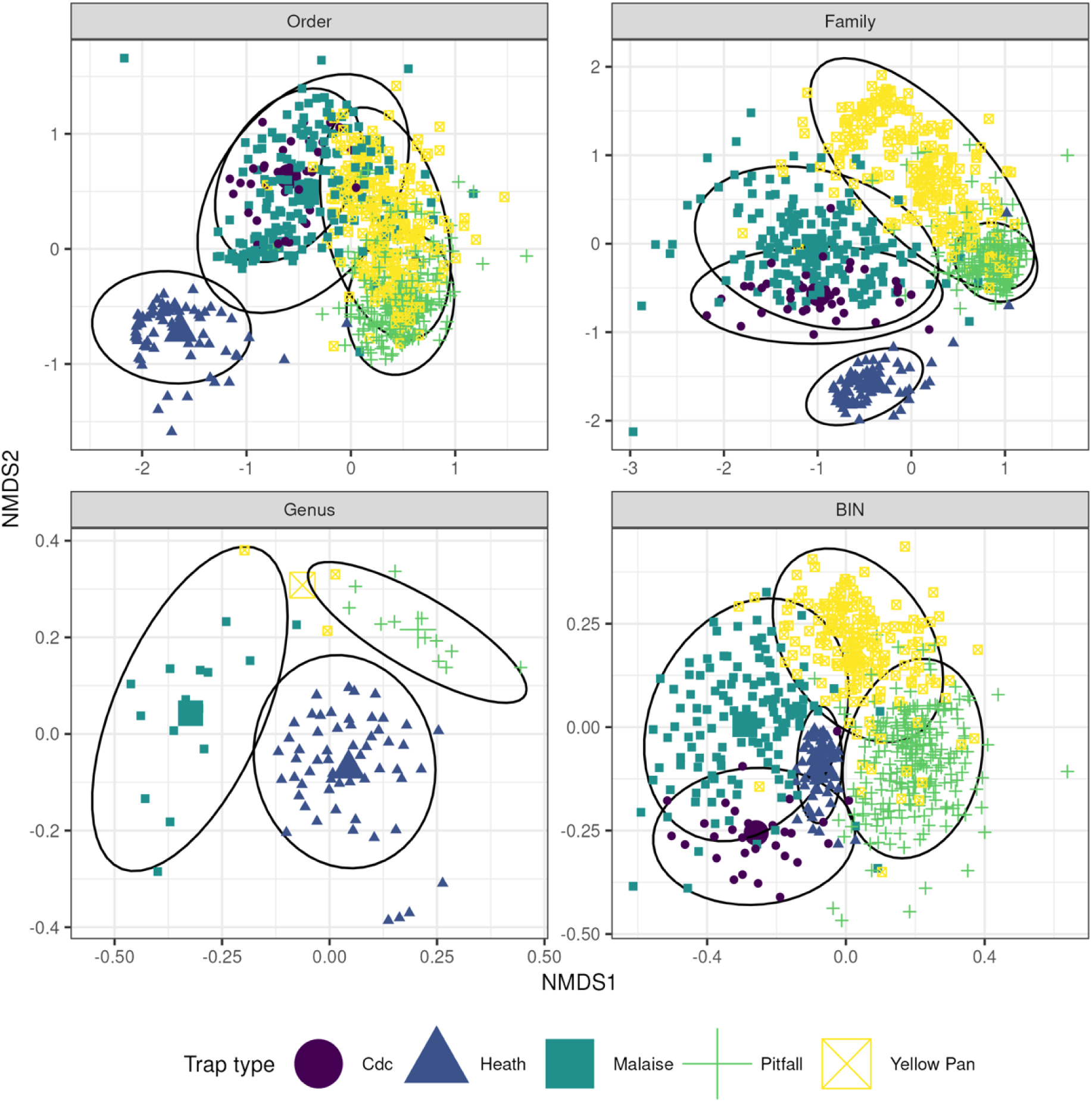
NMDS plots comparing insect assemblages among trap types. Each plot differs in terms of the taxonomic resolution at which samples were categorised. The larger points are the centroids for each trap type. There are fewer data points for the genus-level plot because many samples were not assigned to genus level, despite being identified to family level and assigned a BIN.

Assemblages did not differ significantly among forest, semi-natural, or village habitat types at order (F_4,2_ = 1.3, p = 0.2, R^2^ = 0.002 ) and family (F_4,2_ = 1.4, p = 0.09, R^2^ = 0.002) levels; minor differences emerge at the levels of genus (F_3,2_ = 1.4, p = 0.02, R^2^ = 0.03) and BIN (F_4,2_ = 2.3, p<0.001, R^2^ = 0.006) although the effect sizes and R^2^ indicate trivial effects at most (Supplementary Figure 4).

### Temporal analyses

Malaise traps captured significantly more individual insects during the day (x = 49.1±63.9) than at night (x = 20±25.8; F_1,84_ = 13.5; p < 0.001). This trend was consistent across almost all taxa (Figure 3A), although driven especially by Diptera and Hymenoptera. Many families were substantially more abundant during the day than night (e.g. Chloropidae, Muscidae, Ceratopogonidae, Cecidomyiidae, Formicidae; see Supplementary Table 6), while a few were more abundant at night than during the day (Crambidae, Aphrophoridae, Erebidae, Euteliidae, Gracillariidae). The traps also captured a greater number of unique BINs during the day (x = 32.4±41) than at night (x = 14.6±17.6) (F_1, 84_ = 12.8; p < 0.005) (Figure 3B). The trend of greater variability in captures was significant both at the level of number of insects (F_1,78.7_ = 10.6; p < 0.005) and number of BINs (F_1, 84_ = 9.6; p < 0.005). Ordinations of diurnal and nocturnal assemblages differed slightly at order (F_1,2_ = 9, p<0.001, R^2^=0.06), family (F_1,2_ = 8.4, p<0.001, R^2^=0.06) and BIN (F_1,2_ = 2.4, p<0.01, R^2^ = 0.02) levels but not at the genus level (F_1,2,_ = 1, p = 0.4, R^2^ = 0.07) (see Supplementary Figure 7).

**Figure 3:**
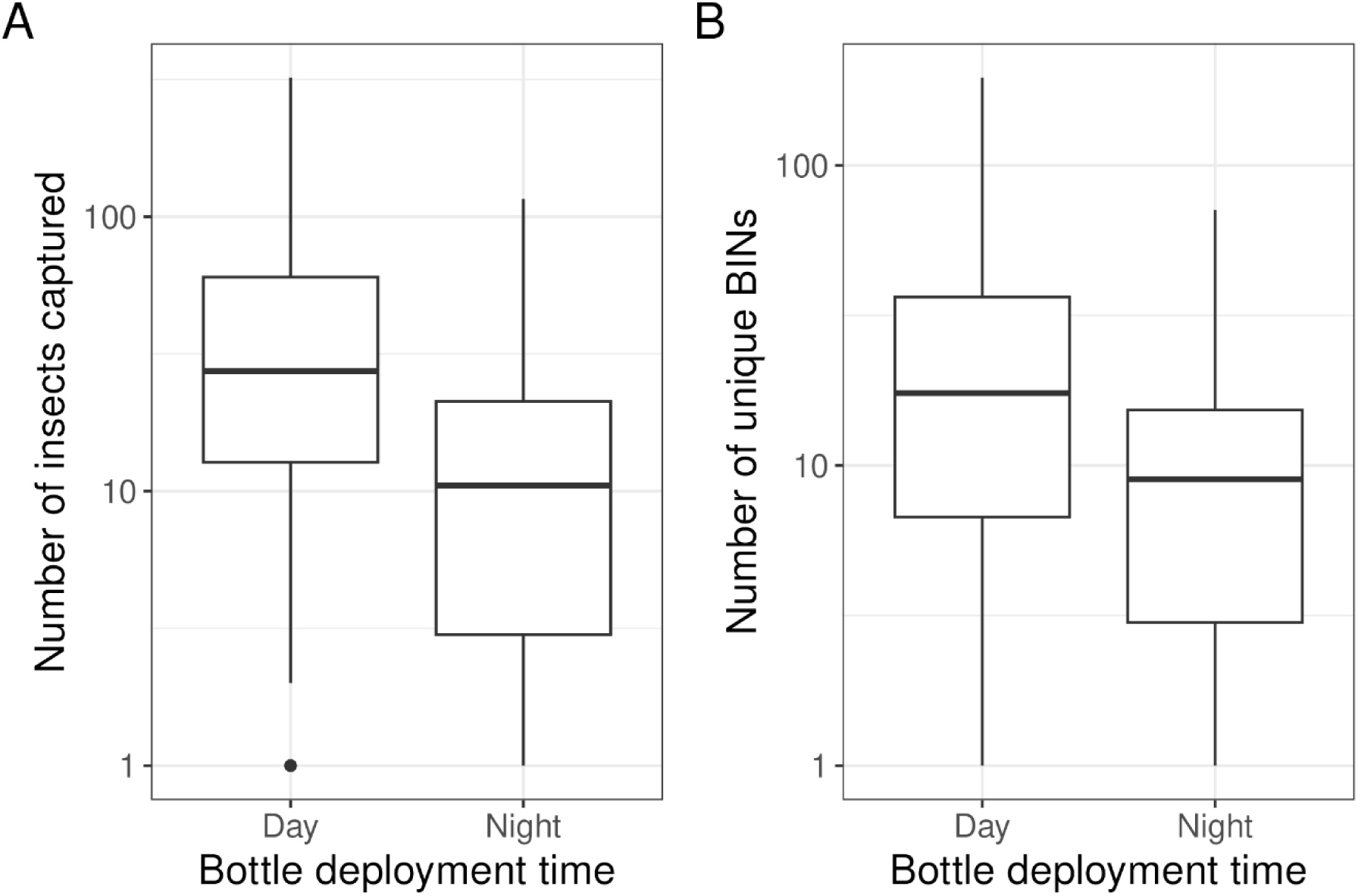
Boxplots showing A) the number of insects captured during diurnal and nocturnal Malaise trap deployment; B) the number of unique BINs captured during diurnal and nocturnal Malaise trap deployment. Y-axes are on a log scale. Boxes show the median, interquartile range and 1.5x the interquartile range.

### Geographic analyses

Of the 10,120 unique BINs recorded in our project, 3,281 (32.3%) were found in the publicly available BOLD dataset. 2,625 BINs (25.9%) were present on BOLD but with no publicly available records (i.e. only found in private datasets). Records of these shared BINs came from 189 countries in seven geographic regions (Table 3). The number of shared BINs detected per taxonomic order was broadly consistent with the abundance of each order in our traps (Table 2, Table 3, Supplementary Table 5): our captures were dominated by Coleoptera, Diptera, Hemiptera, Hymenoptera and Lepidoptera, and these orders also had the most BINs detected in public datasets. Of the 200 most-abundant BINs in our dataset, 104 were already publicly available, 86 had already been sequenced but no other representatives of that BIN were publicly available on BOLD, and 10 were unique to our project. Only eight of the 200 most abundant BINs were assigned binomial names by BOLD (*Carpophilus marginellus*, *Corynoptera forcipata*, *Euplatypus hintzi*, *Hycleus hermanniae*, *Microvelia pygmaea*, *Monolepta jacksoni*, *Nysius graminicola*, and *Peregrinus maidis*). Four of these are considered crop pest species (Supplementary Table 3). BINs were assigned binomial names in 355 cases, accounting for 2.6% of our total samples.

**Table 3:** The number of BINs found in our dataset that have been found in each major geographic region.

| Order name | East Asia | Europe | Latin | Middle | North | South | Sub- | Already | Unique to |
| --- | --- | --- | --- | --- | --- | --- | --- | --- | --- |

|  | **& Pacific** | **&**<br>**Central**<br>**Asia** | **America**<br>**&**<br>**Caribbean** | **East &**<br>**North**<br>**Africa** | **America** | **Asia** | **Saharan**<br>**Africa** | **sequenced,**<br>**no public**<br>**sequences** | **our project** |
| --- | --- | --- | --- | --- | --- | --- | --- | --- | --- |
| Araneae | 2 | 2 | 3 | 1 | 1 | 3 | 25 | 9 | 34 |
| Blattodea | 0 | 0 | 1 | 0 | 0 | 0 | 13 | 19 | 15 |
| Coleoptera | 34 | 23 | 32 | 19 | 16 | 37 | 236 | 532 | 1057 |
| Dermaptera | 0 | 0 | 0 | 1 | 1 | 0 | 2 | 2 | 5 |
| Diptera | 97 | 34 | 62 | 116 | 28 | 116 | 971 | 761 | 935 |
| Entomobryomorpha | 3 | 0 | 3 | 2 | 1 | 2 | 2 | 12 | 11 |
| Ephemeroptera | 0 | 0 | 0 | 1 | 0 | 0 | 1 | 0 | 11 |
| Hemiptera | 46 | 27 | 32 | 50 | 23 | 51 | 309 | 408 | 591 |
| Hymenoptera | 30 | 22 | 24 | 48 | 14 | 24 | 415 | 376 | 854 |
| Lepidoptera | 75 | 81 | 30 | 99 | 32 | 75 | 1027 | 401 | 529 |
| Mantodea | 0 | 0 | 0 | 0 | 0 | 0 | 4 | 1 | 5 |
| Mesostigmata | 0 | 1 | 1 | 1 | 1 | 2 | 2 | 2 | 3 |
| Neuroptera | 0 | 0 | 0 | 1 | 0 | 0 | 7 | 7 | 9 |
| Odonata | 1 | 0 | 1 | 0 | 1 | 1 | 2 | 1 | 0 |
| Orthoptera | 2 | 9 | 1 | 6 | 1 | 3 | 46 | 63 | 61 |
| Plecoptera | 0 | 0 | 0 | 0 | 0 | 0 | 1 | 0 | 1 |
| Poduromorpha | 1 | 0 | 0 | 0 | 0 | 0 | 1 | 0 | 1 |
| Psocodea | 9 | 0 | 13 | 3 | 4 | 9 | 16 | 3 | 6 |
| Thysanoptera | 2 | 1 | 2 | 2 | 1 | 2 | 9 | 14 | 17 |
| Trichoptera | 0 | 0 | 0 | 0 | 0 | 0 | 16 | 3 | 10 |

Thirteen of the top 20 countries sharing the most public BINs with our project are in Africa (Table 4, Figures 4 and 5). In the overall model, the number of shared BINS decreased with geographic distance and increasing number of public BINs (F_2,166_ = 3.76; p = <0.05; R^2^ = 0.03). However when examining the model’s two fixed effects independently, the number of shared BINs correlated positively with the number of samples barcoded in each country (β = 2.12, p<0.05) (Figure 4) but not with the distance between the country and Ghana (β = -1.92, p >0.05) (Figure 5).

**Figure 4:**
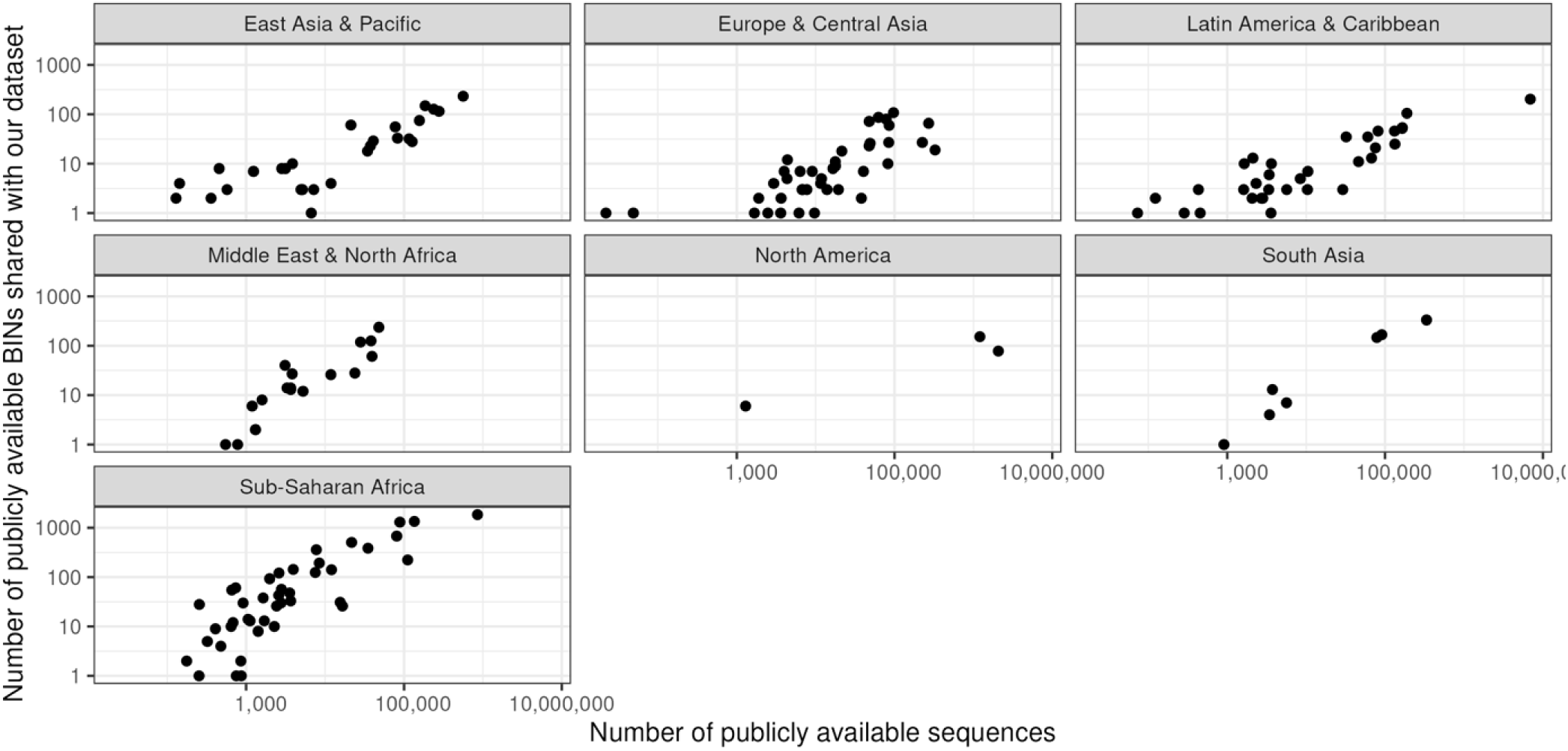
the total number of public sequences available from every country, and the number of BINs that each country shares with our dataset.

**Figure 5:**
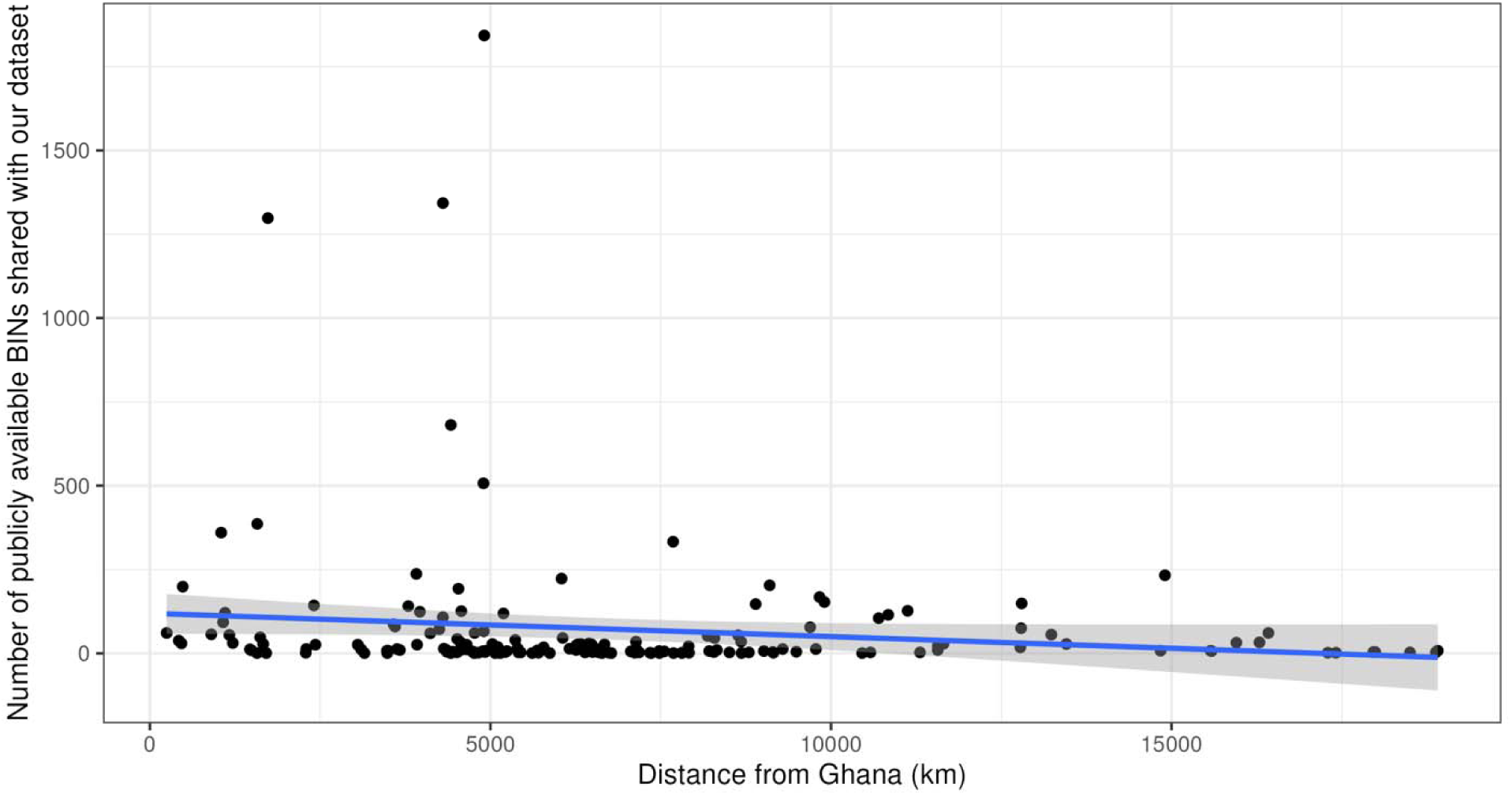
the number of public BINs shared with our dataset per-country, and the shortest distance between that country’s centroid and the centroid of Ghana.

**Table 4:**
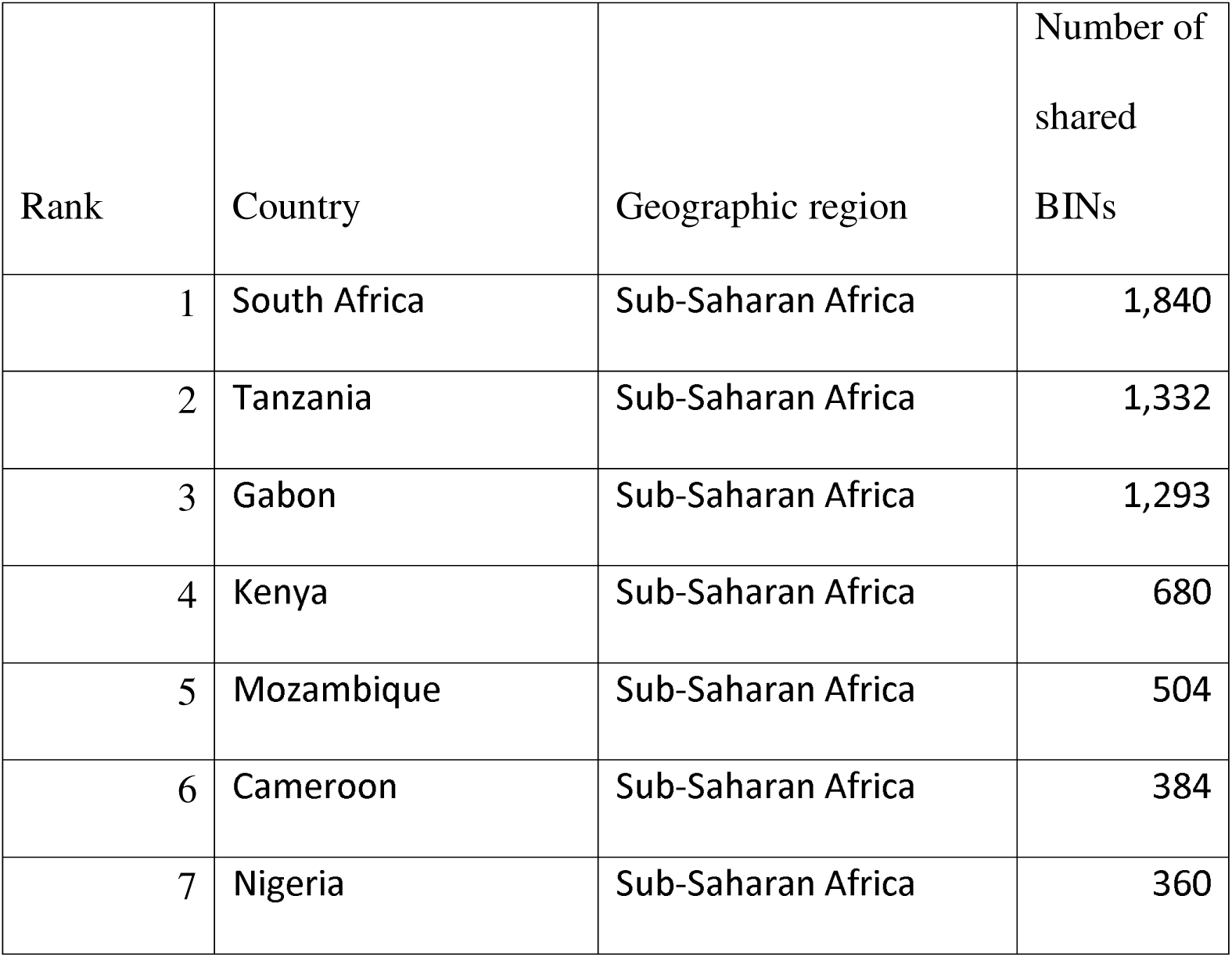
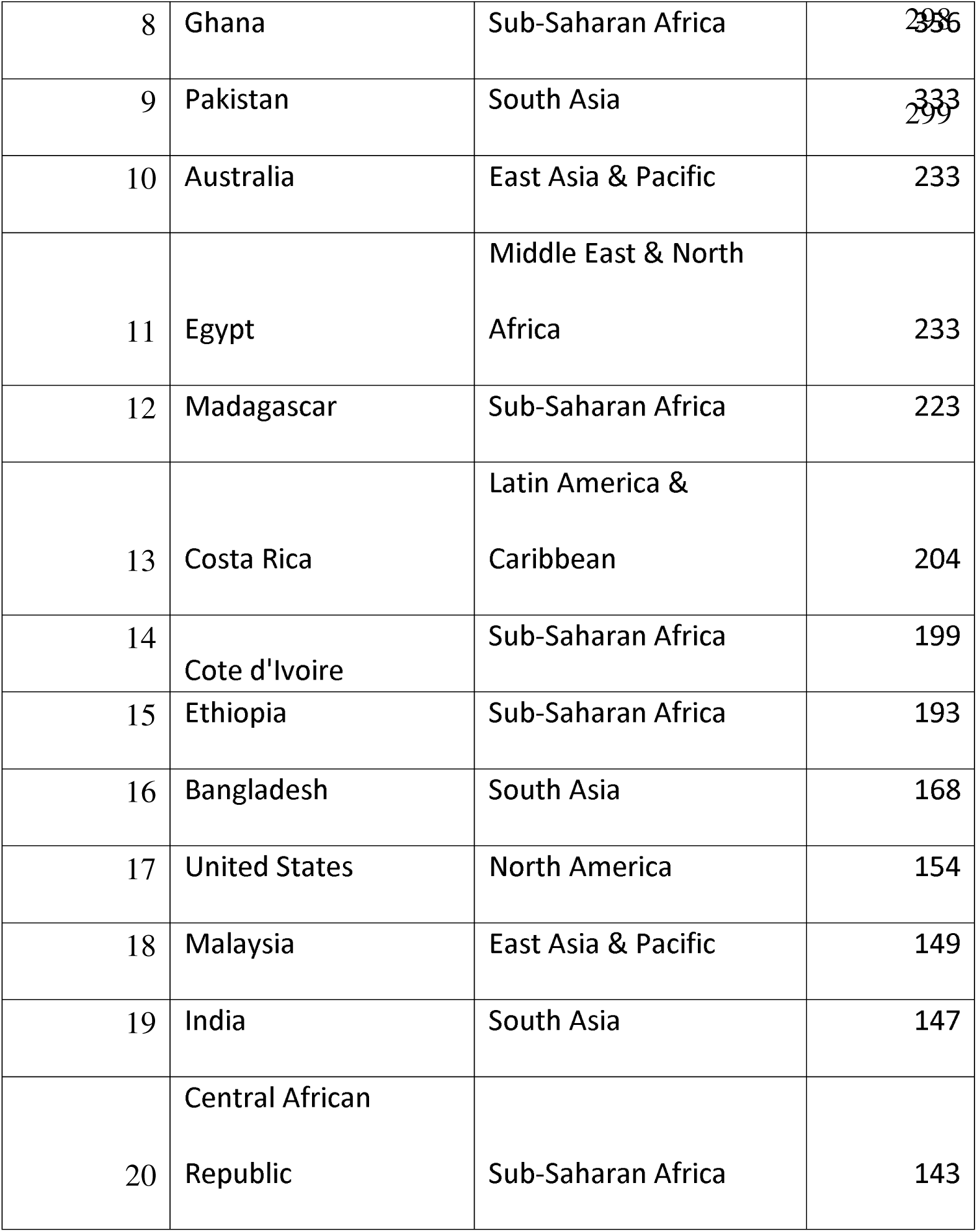
The 20 countries sharing the most publicly available BINs with our dataset

## Discussion

We found over 10,000 unique BINs, more than doubling the number previously documented for West Africa. Despite this intense sampling effort, our species accumulation curves did not asymptote for any taxonomic group or trap type, highlighting the high biodiversity of the sites studied. There was greater overlap in BINs with South Africa than any other country, despite the large geographic distance between the two countries, highlighting the low taxonomic completeness from previous work for much of the region. Our data also show the importance of using multiple trapping methods when documenting arthropod assemblages and highlight the distinct taxonomic groups that are captured by the trap types used, patterns that are especially clear when samples are identified to a fine taxonomic resolution.

### Efficiency and complementarity of trapping methods

Each trap type captured a distinct subset of the arthropod assemblage [32], with our five trap types capturing 60.3% of the estimated total BINs present at the study sites (Supplementary Figure 6). Effective biodiversity survey design requires an understanding of how trap complementarity and overlap influence sampling completeness, allowing effort and cost to be optimised for the question at hand. Generally, Malaise trap samples are consistently dominated by the same 20 insect families, primarily Diptera and Hymenoptera, across continents and biomes [33]. In our study, 14 of these globally dominant families were also among the top 20 most abundant families we recorded in Malaise traps, with five of the remaining six being Diptera (see Supplementary Table 5). While many large-scale insect survey campaigns rely solely on Malaise traps [19–22,34], our results confirm that using a broader range of sampling methods will provide more comprehensive assessments than increased sampling effort with a single trap type [32,35,36]. Determining the species accumulation benefit of any given trap type or combination would require a dedicated study in an area with a well-characterised community (e.g. Wytham Woods, UK [37] or Zackenberg, Greenland [38]) but would be a valuable insight for study design.

Arthropod assemblages captured in pitfall and yellow pan traps had high order-level and family-level overlap, but those captured using other trap types were more distinct. The distinction between assemblages caught by yellow pan and pitfall traps is only clear at the BIN level (Figures 2 and 3, Supplementary Table 2). Both trap types are placed on the ground and so can capture cursorial arthropods. However, yellow pan traps generally target flying, flower-visiting insects that are attracted to the colour yellow [36]. It appears that our yellow pan traps functioned to capture these flower-visitors (overlapping substantially with Malaise traps) as well as many of the cursorial species captured by pitfall traps. At the BIN level, a distinction between the two trap types becomes apparent. Taxa that were much more frequent in yellow pan traps than pitfall traps included Diptera families such as Chloropidae and Sarcophagidae, which are known to be flower visitors. Conversely, Sphaeroceridae were more frequent in pitfall traps than pan traps; many Sphaeroceridae are saprophagous and they may have been attracted to pitfall traps as potential egg laying sites, drawn by the decomposition smell from trapped invertebrates.

While sampling effort (number of insects sequenced) was greatest for Heath traps, the highest rate of BIN accumulation observed was for Malaise traps (Supplementary Figure 6). Heath traps produced the greatest overall number of BINs for the study, and with a distinct subset of the community compared to other trap types, even at the family-level. Heath traps (and other light traps) are commonly employed to target nocturnal Lepidoptera; our results suggest that they may also be an effective way to sample and discover a broad range of other taxa, especially if deployed alongside complementary trapping methods. Budget limitations precluded us from sequencing all trapped arthropods (likely more than 1 million individuals). These results might to some extent be biased by our sub-sorting approach, but we expect the impact to be minimal as sub-sorting was designed to capture the diversity of traps rather than abundance.

### Taxa of potential human importance

A very small fraction of all BINs (0.25%) and individuals (0.6%) corresponded to known crop pest species (Supplementary Table 3). The most common of these species included the Lepidoptera species *Thaumatotibia leucotreta* and Hemiptera species *Nesidiocoris tenuis* and *Rhopalosiphum rufiabdominale*, known pests of peppers and tomatoes, both crops commonly grown at our study sites. Although it is documented to cause significant crop damage in the study areas [39], the important crop pest Fall Armyworm, *Spodoptera frugiperda* (Lepidoptera, Noctuidae) was only detected twice in our data set, and was only captured in Heath traps.

Despite including CO_2_-baited CDC traps specifically deployed to catch blood-feeding Diptera, only 3.1% of all BINS and 1.9% of all individuals trapped were from families that contain blood-feeding species (Supplementary Table 4). We caution that most of these BINs were not identified to species and most families in question (e.g. the especially abundant family Ceratopogonidae) contain both blood-feeding and non-blood-feeding taxa. The three nearly exclusively haematophagous families most likely to be of concern to human health (Culicidae, Simuliidae, and Tabanidae) comprised 0.5% of all captured BINs and 0.28% of all captured insects. The commonest of these families was Ceratopogonidae (mainly caught in Heath traps). Tabanidae were mostly caught in Malaise Traps, and Culicidae were mostly caught in CDC traps (Supplementary Table 4). Taken together, these results show the utility of different trapping methods for surveillance of economically important tropical insects, while highlighting their relatively minor contribution to the overall insect assemblage when compared to the range of numerous economically-neutral or beneficial insects.

### Diurnal activity patterns

In our malaise traps, more insFect individuals and more BINs were trapped during the day than at night. This pattern was driven largely by Diptera and Hymenoptera, (Supplementary Table 6). Surprisingly, inspection of the order and family-level taxonomic composition of Malaise trap samples revealed few other taxa with strongly day-biased or night-biased activity. Activity patterns were more variable during daytime than at night, potentially reflecting higher between-day variance in thermal conditions, which strongly influence insect activity [40] during daylight hours. Taken together, these results suggest that, for passive trapping methods such as Malaise traps, efforts should be made to standardise the extent to which deployments span diurnal and nocturnal periods.

## Conclusions

Our dataset provides insights into our current knowledge of tropical arthropod biodiversity and highlights the need for extensive further research in the region and beyond. By publishing this manuscript and dataset we provide a genetic and taxonomic resource for the scientific community, particularly for those studying tropical arthropod fauna in West Africa. While BINs are an invaluable first step in describing biodiversity, we encourage efforts towards the formal taxonomic description of the many unnamed taxa within this understudied arthropod assemblage.

## Authors’ contributions

DRHB: methodology, formal analysis, data curation, writing (original draft and review and editing), visualisation ED: investigation, resources, data curation, writing (review and editing) BA: investigation, resources, writing (review and editing) NA: investigation, resources, writing (review and editing) EDO: investigation, resources, writing (review and editing) SAA: investigation, resources, writing (review and editing) AB: investigation, resources, writing (review and editing) HCJG: conceptualisation, methodology, writing (review and editing), supervision, project administration, funding acquisition OTL: conceptualisation, methodology, writing (review and editing), supervision, project administration, funding acquisition FAA: conceptualisation, methodology, writing (review and editing), supervision, project administration, funding acquisition TDH: conceptualisation, methodology, formal analysis, investigation, data curation, writing (review and editing), supervision, project administration

## Acknowledgements

This research was supported by members of the Target Malaria Ghana Stakeholder Engagement Team, especially Divine Dzokoto and Linda Mawutor Aboagye. Data collection was facilitated by village elders, members of the two study communities, and especially those who assisted with trap deployment and sample collections. Sample sorting protocols and DNA barcoding processing was supported by the Centre for Biodiversity Genomics, especially Michelle D’Souza and Jayme Sones. CDC trap samples were sorted by Ben Bellekom. All data are stored on Earthcape Biodiversity Database Platform (https://earthcape.com/) with the support of Evgeniy Meyke. All authors were working as members of the Target Malaria Not-for-Profit Research Consortium (www.targetmalaria.org), which receives core funding from the Gates Foundation and from Open Philanthropy.

