## Supplementary information for "Characterising a species-rich and understudied tropical insect fauna using DNA barcoding"

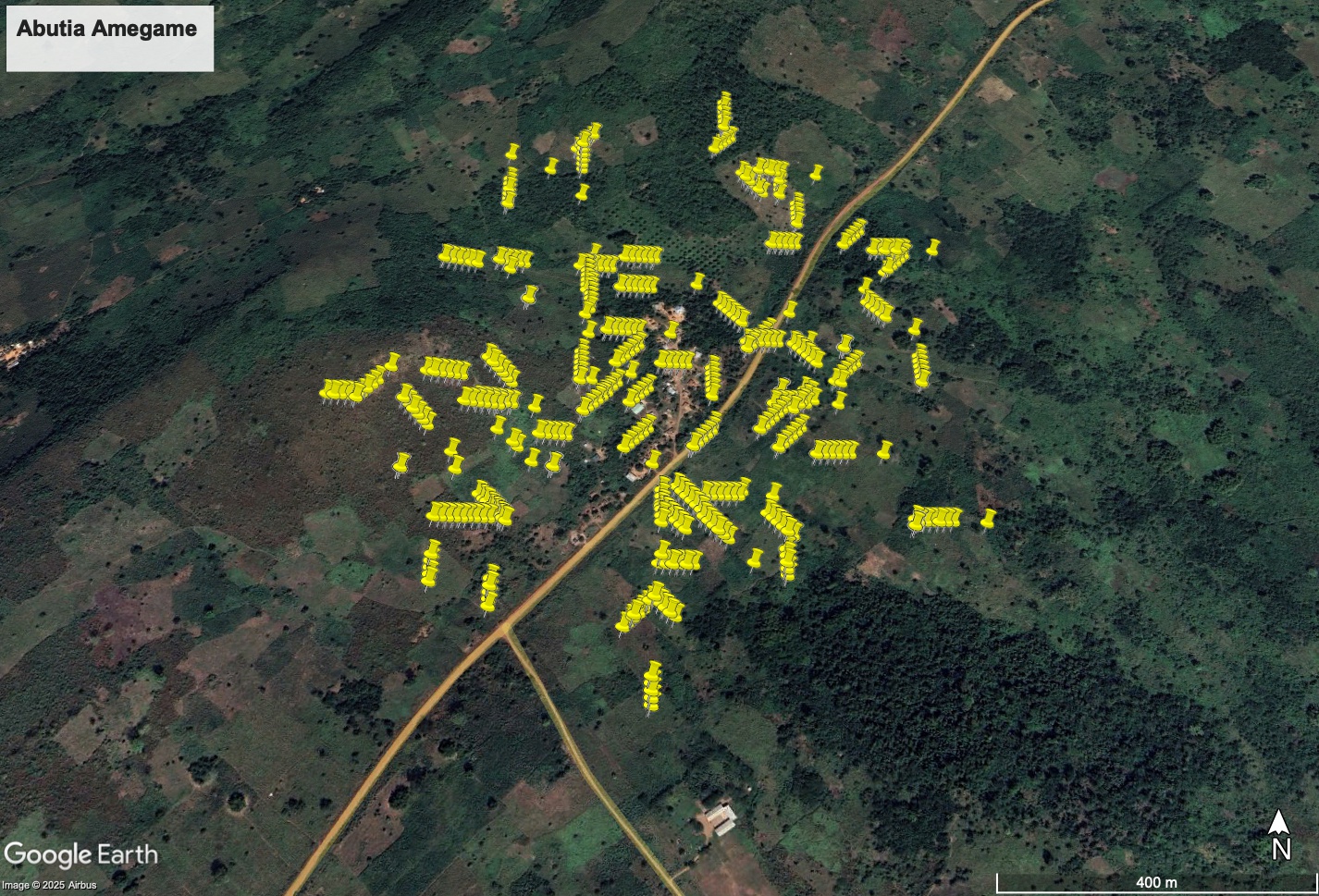

Supplementary figure 1: map of random sampling points at the village Abutia Amegame.

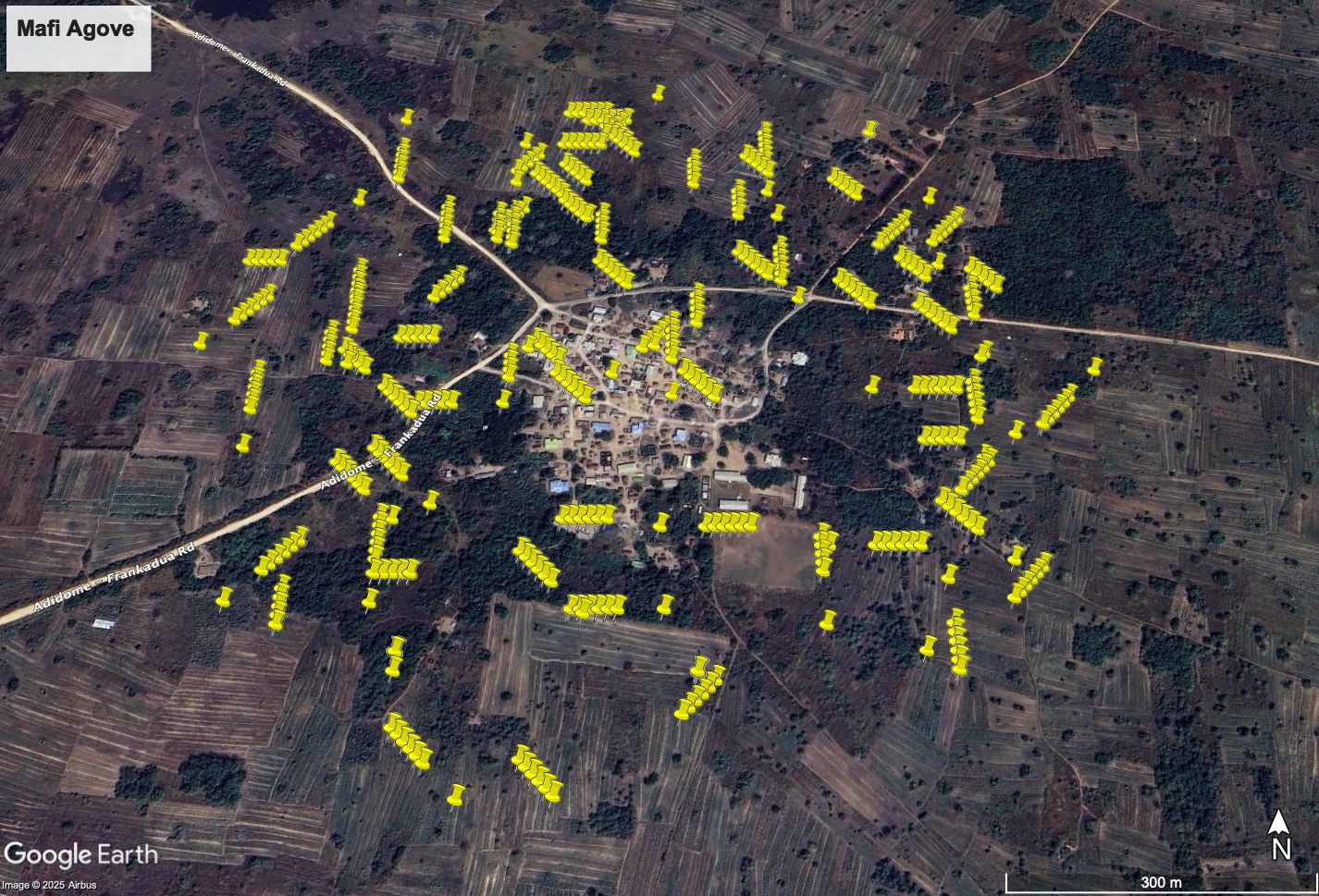

Supplementary figure 2:map of random sampling points at the village Mafi Agove.

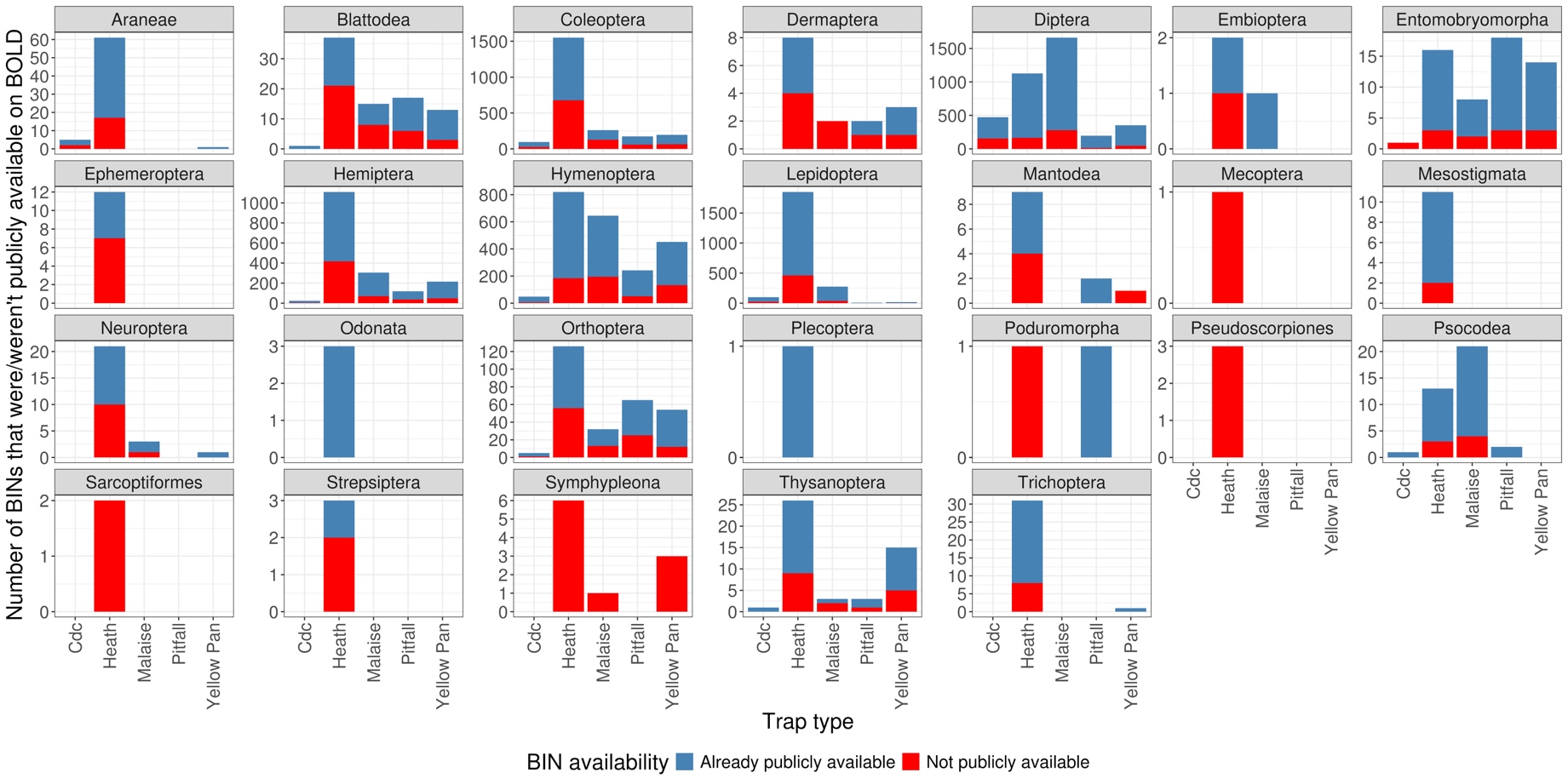

Supplementary Figure 3: Public availability of BINs detected, by taxonomic order and trap-type.

Supplementary Table 1: The number of BINs of a given order sequenced in each trap type.

| **Order** | **CDC** | **Heath** | **Malaise** | **Pitfall** | **Yellow Pan** |
| --- | --- | --- | --- | --- | --- |
| Araneae | 15 | 140 | 0 | 0 | 2 |
| Archaeognatha | 0 | 3 | 0 | 4 | 4 |
| Blattodea | 2 | 389 | 39 | 104 | 58 |
| Coleoptera | 408 | 21560 | 671 | 1111 | 613 |
| Dermaptera | 2 | 75 | 5 | 11 | 6 |
| Diptera | 1758 | 6140 | 6550 | 599 | 2078 |
| Embioptera | 0 | 8 | 1 | 0 | 0 |
| Entomobryomorpha | 2 | 60 | 36 | 227 | 176 |
| Ephemeroptera | 0 | 215 | 0 | 0 | 0 |
| Hemiptera | 140 | 12786 | 1363 | 465 | 1081 |
| Hymenoptera | 305 | 6215 | 1993 | 4851 | 2654 |
| Isopoda | 0 | 1 | 0 | 2 | 0 |
| Ixodida | 0 | 0 | 0 | 3 | 0 |
| Lepidoptera | 340 | 14768 | 817 | 18 | 41 |
| Mantodea | 1 | 37 | 0 | 3 | 4 |
| Mecoptera | 0 | 16 | 0 | 0 | 0 |
| Megaloptera | 0 | 1 | 0 | 0 | 0 |
| Mesostigmata | 0 | 59 | 0 | 0 | 0 |
| Neuroptera | 0 | 112 | 19 | 4 | 2 |
| Odonata | 2 | 4 | 3 | 0 | 2 |
| Orthoptera | 13 | 1762 | 220 | 829 | 389 |
| Phasmida | 0 | 0 | 0 | 1 | 0 |
| Plecoptera | 3 | 3 | 1 | 1 | 1 |
| Poduromorpha | 0 | 1 | 0 | 1 | 0 |
| Pseudoscorpiones | 0 | 5 | 0 | 0 | 0 |
| Psocodea | 18 | 82 | 98 | 8 | 19 |
| Sarcoptiformes | 0 | 2 | 0 | 0 | 0 |
| Strepsiptera | 0 | 19 | 0 | 0 | 0 |
| Symphypleona | 0 | 8 | 3 | 0 | 3 |
| Thysanoptera | 2 | 135 | 4 | 8 | 54 |
| Trichoptera | 0 | 454 | 113 | 1 | 3 |

Supplementary Table 2: The number of BINs of a given family sequenced in each trap type.

| **Order** | **Family** | **CDC** | **Heath** | **Malaise** | **Pitfall** | **Yellow Pan** |
| --- | --- | --- | --- | --- | --- | --- |
| Araneae | Araneidae | 2 | 9 | 0 | 0 | 0 |
| Araneae | Cheiracanthiidae | 0 | 42 | 0 | 0 | 0 |
| Araneae | Clubionidae | 0 | 2 | 0 | 0 | 0 |
| Araneae | Ctenidae | 0 | 1 | 0 | 0 | 0 |
| Araneae | Dictynidae | 0 | 2 | 0 | 0 | 0 |
| Araneae | Gnaphosidae | 0 | 1 | 0 | 0 | 0 |
| Araneae | Linyphiidae | 2 | 4 | 0 | 0 | 0 |
| Araneae | Lycosidae | 0 | 11 | 0 | 0 | 0 |
| Araneae | Mysmenidae | 1 | 1 | 0 | 0 | 0 |
| Araneae | Oxyopidae | 0 | 8 | 0 | 0 | 0 |
| Araneae | Philodromidae | 0 | 1 | 0 | 0 | 0 |
| Araneae | Pholcidae | 0 | 1 | 0 | 0 | 0 |
| Araneae | Pisauridae | 0 | 6 | 0 | 0 | 0 |
| Araneae | Salticidae | 0 | 2 | 0 | 0 | 1 |
| Araneae | Scytodidae | 0 | 1 | 0 | 0 | 0 |
| Araneae | Tetragnathidae | 0 | 2 | 0 | 0 | 0 |
| Araneae | Theridiidae | 0 | 21 | 0 | 0 | 0 |
| Araneae | Thomisidae | 1 | 11 | 0 | 0 | 0 |
| Araneae | Trachelidae | 0 | 2 | 0 | 0 | 0 |
| Araneae | Zodariidae | 0 | 2 | 0 | 0 | 0 |
| Blattodea | Blaberidae | 0 | 1 | 0 | 0 | 0 |
| Blattodea | Blattellidae | 1 | 50 | 0 | 1 | 1 |
| Blattodea | Blattidae | 0 | 0 | 0 | 2 | 1 |
| Blattodea | Ectobiidae | 0 | 18 | 3 | 1 | 2 |
| Blattodea | Kalotermitidae | 0 | 10 | 0 | 0 | 0 |
| Blattodea | Rhinotermitidae | 0 | 14 | 0 | 0 | 0 |
| Blattodea | Termitidae | 0 | 132 | 18 | 58 | 22 |
| Coleoptera | Acanthocnemidae | 0 | 3 | 0 | 0 | 0 |
| Coleoptera | Aderidae | 3 | 35 | 16 | 1 | 2 |
| Coleoptera | Anthicidae | 1 | 94 | 20 | 59 | 24 |
| Coleoptera | Anthribidae | 0 | 5 | 20 | 0 | 0 |
| Coleoptera | Archeocrypticidae | 0 | 8 | 0 | 0 | 0 |
| Coleoptera | Attelabidae | 0 | 2 | 2 | 0 | 2 |
| Coleoptera | Biphyllidae | 2 | 10 | 1 | 0 | 3 |
| Coleoptera | Bostrichidae | 2 | 184 | 9 | 0 | 0 |
| Coleoptera | Bothrideridae | 0 | 1 | 0 | 0 | 0 |
| Coleoptera | Brentidae | 1 | 69 | 3 | 0 | 3 |
| Coleoptera | Buprestidae | 0 | 0 | 8 | 13 | 4 |
| Coleoptera | Cantharidae | 0 | 5 | 1 | 0 | 0 |
| Coleoptera | Carabidae | 9 | 2787 | 11 | 108 | 22 |
| Coleoptera | Cerambycidae | 0 | 75 | 7 | 0 | 0 |
| Coleoptera | Cerylonidae | 1 | 9 | 0 | 0 | 0 |
| Coleoptera | Chrysomelidae | 22 | 1097 | 118 | 77 | 162 |
| Coleoptera | Ciidae | 0 | 8 | 0 | 0 | 0 |
| Coleoptera | Cleridae | 0 | 9 | 6 | 0 | 2 |
| Coleoptera | Coccinellidae | 4 | 40 | 32 | 2 | 23 |
| Coleoptera | Corylophidae | 0 | 21 | 10 | 0 | 1 |
| Coleoptera | Cryptophagidae | 0 | 20 | 0 | 0 | 0 |
| Coleoptera | Curculionidae | 15 | 523 | 19 | 2 | 8 |
| Coleoptera | Cybocephalidae | 0 | 0 | 2 | 0 | 0 |
| Coleoptera | Dermestidae | 0 | 17 | 0 | 0 | 0 |
| Coleoptera | Dytiscidae | 1 | 1219 | 0 | 0 | 1 |
| Coleoptera | Elateridae | 2 | 380 | 14 | 10 | 5 |
| Coleoptera | Elmidae | 0 | 17 | 0 | 0 | 0 |
| Coleoptera | Endomychidae | 0 | 4 | 0 | 1 | 0 |
| Coleoptera | Erotylidae | 2 | 95 | 3 | 1 | 3 |
| Coleoptera | Eucinetidae | 0 | 1 | 0 | 0 | 0 |
| Coleoptera | Euxestidae | 0 | 23 | 0 | 0 | 0 |
| Coleoptera | Geotrupidae | 0 | 2 | 0 | 0 | 0 |
| Coleoptera | Gyrinidae | 0 | 7 | 0 | 0 | 0 |
| Coleoptera | Haliplidae | 0 | 1 | 0 | 0 | 0 |
| Coleoptera | Heteroceridae | 0 | 365 | 0 | 0 | 0 |
| Coleoptera | Histeridae | 0 | 3 | 0 | 14 | 0 |
| Coleoptera | Hybosoridae | 0 | 51 | 0 | 0 | 1 |
| Coleoptera | Hydraenidae | 0 | 27 | 0 | 0 | 0 |
| Coleoptera | Hydrophilidae | 1 | 1771 | 0 | 2 | 2 |
| Coleoptera | Laemophloeidae | 0 | 27 | 3 | 0 | 0 |
| Coleoptera | Lampyridae | 0 | 18 | 0 | 0 | 0 |
| Coleoptera | Latridiidae | 7 | 28 | 3 | 0 | 7 |
| Coleoptera | Leiodidae | 0 | 10 | 0 | 0 | 0 |
| Coleoptera | Limnichidae | 0 | 177 | 0 | 0 | 0 |
| Coleoptera | Lycidae | 0 | 22 | 2 | 1 | 0 |
| Coleoptera | Lymexylidae | 0 | 3 | 0 | 0 | 0 |
| Coleoptera | Meloidae | 1 | 11 | 4 | 187 | 53 |
| Coleoptera | Melyridae | 0 | 0 | 4 | 8 | 9 |
| Coleoptera | Monotomidae | 0 | 0 | 0 | 1 | 0 |
| Coleoptera | Mordellidae | 5 | 73 | 36 | 0 | 6 |
| Coleoptera | Mycetophagidae | 0 | 15 | 1 | 2 | 1 |
| Coleoptera | Mycteridae | 0 | 5 | 2 | 0 | 0 |
| Coleoptera | Nitidulidae | 0 | 239 | 11 | 72 | 13 |
| Coleoptera | Noteridae | 0 | 165 | 0 | 0 | 0 |
| Coleoptera | Oedemeridae | 0 | 1 | 0 | 0 | 0 |
| Coleoptera | Passandridae | 0 | 1 | 0 | 0 | 0 |
| Coleoptera | Phalacridae | 46 | 156 | 36 | 5 | 18 |
| Coleoptera | Psephenidae | 0 | 0 | 0 | 1 | 0 |
| Coleoptera | Ptiliidae | 0 | 3 | 1 | 0 | 0 |
| Coleoptera | Ptinidae | 0 | 17 | 8 | 0 | 0 |
| Coleoptera | Salpingidae | 0 | 2 | 0 | 0 | 0 |
| Coleoptera | Scarabaeidae | 16 | 1317 | 8 | 10 | 13 |
| Coleoptera | Scirtidae | 0 | 86 | 2 | 0 | 0 |
| Coleoptera | Scraptiidae | 0 | 10 | 8 | 0 | 1 |
| Coleoptera | Silvanidae | 0 | 26 | 0 | 0 | 1 |
| Coleoptera | Spercheidae | 0 | 32 | 0 | 0 | 0 |
| Coleoptera | Sphindidae | 0 | 1 | 0 | 0 | 1 |
| Coleoptera | Staphylinidae | 26 | 4508 | 16 | 93 | 43 |
| Coleoptera | Tenebrionidae | 2 | 86 | 3 | 19 | 0 |
| Coleoptera | Throscidae | 0 | 3 | 9 | 0 | 0 |
| Coleoptera | Zopheridae | 0 | 15 | 0 | 0 | 0 |
| Dermaptera | Forficulidae | 0 | 3 | 0 | 2 | 2 |
| Dermaptera | Labiduridae | 0 | 0 | 0 | 4 | 1 |
| Dermaptera | Spongiphoridae | 0 | 47 | 1 | 2 | 3 |
| Diptera | Agromyzidae | 1 | 3 | 77 | 0 | 3 |
| Diptera | Anthomyiidae | 1 | 1 | 8 | 1 | 0 |
| Diptera | Anthomyzidae | 0 | 0 | 4 | 0 | 0 |
| Diptera | Asilidae | 0 | 9 | 0 | 0 | 0 |
| Diptera | Asteiidae | 0 | 2 | 3 | 0 | 0 |
| Diptera | Bombyliidae | 0 | 1 | 11 | 2 | 3 |
| Diptera | Calliphoridae | 17 | 66 | 147 | 8 | 21 |
| Diptera | Canacidae | 1 | 1 | 0 | 0 | 0 |
| Diptera | Carnidae | 0 | 0 | 2 | 0 | 0 |
| Diptera | Cecidomyiidae | 805 | 917 | 991 | 47 | 63 |
| Diptera | Celyphidae | 0 | 0 | 2 | 0 | 0 |
| Diptera | Ceratopogonidae | 57 | 1037 | 442 | 1 | 19 |
| Diptera | Chamaemyiidae | 0 | 0 | 3 | 0 | 0 |
| Diptera | Chaoboridae | 0 | 81 | 0 | 0 | 0 |
| Diptera | Chironomidae | 4 | 573 | 145 | 2 | 6 |
| Diptera | Chloropidae | 4 | 210 | 827 | 42 | 142 |
| Diptera | Chyromyidae | 0 | 15 | 19 | 0 | 0 |
| Diptera | Clusiidae | 0 | 0 | 1 | 0 | 0 |
| Diptera | Cryptochetidae | 0 | 0 | 6 | 0 | 0 |
| Diptera | Culicidae | 180 | 6 | 4 | 1 | 1 |
| Diptera | Curtonotidae | 1 | 3 | 0 | 0 | 0 |
| Diptera | Diopsidae | 0 | 6 | 1 | 2 | 6 |
| Diptera | Dolichopodidae | 5 | 123 | 139 | 68 | 1055 |
| Diptera | Drosophilidae | 28 | 50 | 180 | 15 | 21 |
| Diptera | Ephydridae | 5 | 175 | 44 | 51 | 30 |
| Diptera | Glossinidae | 0 | 0 | 2 | 0 | 0 |
| Diptera | Hybotidae | 3 | 36 | 87 | 3 | 29 |
| Diptera | Keroplatidae | 2 | 7 | 2 | 1 | 0 |
| Diptera | Lauxaniidae | 0 | 8 | 47 | 2 | 3 |
| Diptera | Limoniidae | 33 | 117 | 8 | 0 | 1 |
| Diptera | Lonchaeidae | 0 | 14 | 208 | 2 | 1 |
| Diptera | Micropezidae | 0 | 0 | 5 | 0 | 0 |
| Diptera | Milichiidae | 6 | 160 | 431 | 19 | 39 |
| Diptera | Muscidae | 30 | 123 | 659 | 30 | 26 |
| Diptera | Mycetophilidae | 0 | 36 | 21 | 1 | 2 |
| Diptera | Neriidae | 0 | 1 | 6 | 0 | 0 |
| Diptera | Odiniidae | 0 | 1 | 0 | 0 | 0 |
| Diptera | Periscelididae | 0 | 5 | 3 | 0 | 2 |
| Diptera | Phoridae | 7 | 214 | 313 | 97 | 96 |
| Diptera | Pipunculidae | 0 | 3 | 12 | 0 | 17 |
| Diptera | Platypezidae | 0 | 1 | 0 | 0 | 0 |
| Diptera | Platystomatidae | 1 | 12 | 77 | 0 | 1 |
| Diptera | Psychodidae | 25 | 22 | 169 | 7 | 10 |
| Diptera | Pyrgotidae | 0 | 12 | 6 | 0 | 0 |
| Diptera | Sarcophagidae | 7 | 26 | 171 | 10 | 172 |
| Diptera | Scatopsidae | 0 | 6 | 2 | 2 | 0 |
| Diptera | Scenopinidae | 0 | 0 | 2 | 0 | 0 |
| Diptera | Sciaridae | 24 | 837 | 193 | 18 | 26 |
| Diptera | Sciomyzidae | 0 | 1 | 0 | 0 | 0 |
| Diptera | Sepsidae | 0 | 3 | 1 | 0 | 0 |
| Diptera | Simuliidae | 0 | 10 | 2 | 0 | 0 |
| Diptera | Sphaeroceridae | 5 | 467 | 106 | 84 | 17 |
| Diptera | Stratiomyidae | 1 | 23 | 53 | 0 | 10 |
| Diptera | Syrphidae | 4 | 11 | 29 | 3 | 16 |
| Diptera | Tabanidae | 3 | 36 | 21 | 0 | 0 |
| Diptera | Tachinidae | 2 | 34 | 105 | 10 | 10 |
| Diptera | Tephritidae | 1 | 9 | 8 | 0 | 0 |
| Diptera | Ulidiidae | 0 | 53 | 74 | 0 | 11 |
| Embioptera | Oligotomidae | 0 | 7 | 1 | 0 | 0 |
| Entomobryomorpha | Entomobryidae | 0 | 9 | 26 | 5 | 7 |
| Entomobryomorpha | Isotomidae | 0 | 11 | 0 | 135 | 77 |
| Entomobryomorpha | Lepidocyrtidae | 0 | 1 | 0 | 0 | 0 |
| Entomobryomorpha | Seiridae | 0 | 11 | 0 | 0 | 0 |
| Ephemeroptera | Baetidae | 0 | 26 | 0 | 0 | 0 |
| Ephemeroptera | Tricorythidae | 0 | 2 | 0 | 0 | 0 |
| Hemiptera | Achilidae | 3 | 154 | 0 | 0 | 0 |
| Hemiptera | Aleyrodidae | 0 | 55 | 18 | 1 | 11 |
| Hemiptera | Alydidae | 0 | 42 | 1 | 7 | 5 |
| Hemiptera | Anthocoridae | 0 | 29 | 2 | 12 | 6 |
| Hemiptera | Aphididae | 0 | 25 | 33 | 4 | 97 |
| Hemiptera | Aphrophoridae | 0 | 48 | 6 | 1 | 5 |
| Hemiptera | Aradidae | 0 | 34 | 0 | 0 | 0 |
| Hemiptera | Belostomatidae | 0 | 82 | 0 | 0 | 0 |
| Hemiptera | Berytidae | 0 | 7 | 1 | 0 | 0 |
| Hemiptera | Blissidae | 0 | 5 | 0 | 0 | 0 |
| Hemiptera | Caliscelidae | 0 | 0 | 0 | 0 | 1 |
| Hemiptera | Carsidaridae | 0 | 1 | 1 | 0 | 2 |
| Hemiptera | Ceratocombidae | 0 | 239 | 1 | 1 | 3 |
| Hemiptera | Cercopidae | 0 | 5 | 1 | 0 | 0 |
| Hemiptera | Cicadellidae | 40 | 3395 | 774 | 162 | 369 |
| Hemiptera | Cixiidae | 0 | 106 | 2 | 0 | 6 |
| Hemiptera | Coreidae | 0 | 14 | 0 | 1 | 1 |
| Hemiptera | Corixidae | 0 | 181 | 0 | 0 | 0 |
| Hemiptera | Cydnidae | 0 | 311 | 0 | 5 | 0 |
| Hemiptera | Cymidae | 0 | 21 | 0 | 0 | 0 |
| Hemiptera | Delphacidae | 1 | 1515 | 9 | 10 | 73 |
| Hemiptera | Derbidae | 0 | 16 | 2 | 0 | 1 |
| Hemiptera | Diaspididae | 0 | 1 | 0 | 0 | 0 |
| Hemiptera | Dictyopharidae | 0 | 161 | 0 | 2 | 5 |
| Hemiptera | Enicocephalidae | 0 | 2 | 0 | 1 | 0 |
| Hemiptera | Flatidae | 0 | 101 | 3 | 0 | 0 |
| Hemiptera | Fulgoridae | 0 | 29 | 0 | 0 | 0 |
| Hemiptera | Fulgoroidea_incertae_sedis | 0 | 7 | 0 | 1 | 2 |
| Hemiptera | Geocoridae | 0 | 0 | 0 | 0 | 2 |
| Hemiptera | Gerridae | 0 | 22 | 0 | 0 | 0 |
| Hemiptera | Hebridae | 0 | 18 | 0 | 0 | 0 |
| Hemiptera | Heterogastridae | 0 | 1 | 0 | 0 | 0 |
| Hemiptera | Hydrometridae | 0 | 41 | 0 | 0 | 0 |
| Hemiptera | Issidae | 0 | 0 | 5 | 0 | 5 |
| Hemiptera | Kinnaridae | 0 | 1 | 0 | 0 | 0 |
| Hemiptera | Largidae | 0 | 47 | 1 | 0 | 0 |
| Hemiptera | Lasiochilidae | 0 | 2 | 0 | 2 | 0 |
| Hemiptera | Lethaeidae | 0 | 22 | 0 | 0 | 0 |
| Hemiptera | Lophopidae | 0 | 6 | 0 | 0 | 0 |
| Hemiptera | Lygaeidae | 0 | 130 | 10 | 5 | 1 |
| Hemiptera | Machaerotidae | 0 | 1 | 0 | 0 | 0 |
| Hemiptera | Meenoplidae | 0 | 37 | 33 | 0 | 2 |
| Hemiptera | Membracidae | 0 | 4 | 0 | 0 | 3 |
| Hemiptera | Mesoveliidae | 0 | 113 | 0 | 0 | 0 |
| Hemiptera | Miridae | 5 | 519 | 21 | 1 | 4 |
| Hemiptera | Nabidae | 0 | 75 | 0 | 0 | 1 |
| Hemiptera | Ninidae | 0 | 9 | 0 | 0 | 0 |
| Hemiptera | Notonectidae | 0 | 23 | 0 | 0 | 0 |
| Hemiptera | Ochteridae | 0 | 21 | 0 | 0 | 0 |
| Hemiptera | Oxycarenidae | 0 | 1 | 1 | 0 | 0 |
| Hemiptera | Pentatomidae | 0 | 176 | 0 | 1 | 2 |
| Hemiptera | Piesmatidae | 0 | 0 | 0 | 1 | 1 |
| Hemiptera | Plataspidae | 0 | 0 | 0 | 1 | 0 |
| Hemiptera | Pleidae | 0 | 115 | 0 | 0 | 0 |
| Hemiptera | Psyllidae | 0 | 10 | 16 | 3 | 0 |
| Hemiptera | Psylloidea_incertae_sedis | 0 | 5 | 5 | 2 | 6 |
| Hemiptera | Pyrrhocoridae | 0 | 254 | 7 | 16 | 5 |
| Hemiptera | Reduviidae | 0 | 84 | 3 | 8 | 0 |
| Hemiptera | Rhopalidae | 0 | 2 | 0 | 2 | 1 |
| Hemiptera | Rhyparochromidae | 0 | 1125 | 8 | 25 | 13 |
| Hemiptera | Ricaniidae | 0 | 3 | 0 | 0 | 0 |
| Hemiptera | Schizopteridae | 0 | 41 | 3 | 3 | 7 |
| Hemiptera | Stenocephalidae | 0 | 1 | 0 | 0 | 0 |
| Hemiptera | Tettigometridae | 0 | 1 | 0 | 1 | 1 |
| Hemiptera | Tingidae | 0 | 78 | 0 | 5 | 9 |
| Hemiptera | Triozidae | 0 | 0 | 2 | 0 | 2 |
| Hemiptera | Tropiduchidae | 0 | 19 | 0 | 0 | 0 |
| Hemiptera | Veliidae | 0 | 85 | 0 | 2 | 0 |
| Hymenoptera | Agaonidae | 0 | 81 | 0 | 0 | 0 |
| Hymenoptera | Ammoplanidae | 0 | 0 | 1 | 0 | 0 |
| Hymenoptera | Ampulicidae | 0 | 0 | 4 | 0 | 2 |
| Hymenoptera | Aphelinidae | 0 | 5 | 0 | 0 | 2 |
| Hymenoptera | Apidae | 0 | 15 | 21 | 6 | 8 |
| Hymenoptera | Bembicidae | 0 | 0 | 2 | 0 | 0 |
| Hymenoptera | Bethylidae | 3 | 61 | 74 | 22 | 153 |
| Hymenoptera | Braconidae | 9 | 584 | 168 | 3 | 28 |
| Hymenoptera | Ceraphronidae | 0 | 0 | 1 | 0 | 6 |
| Hymenoptera | Chalcididae | 0 | 0 | 12 | 0 | 5 |
| Hymenoptera | Chrysididae | 0 | 0 | 32 | 1 | 3 |
| Hymenoptera | Cleonymidae | 0 | 0 | 1 | 0 | 0 |
| Hymenoptera | Crabronidae | 0 | 2 | 112 | 14 | 307 |
| Hymenoptera | Diapriidae | 4 | 54 | 6 | 5 | 54 |
| Hymenoptera | Dryinidae | 0 | 50 | 8 | 1 | 6 |
| Hymenoptera | Encyrtidae | 0 | 7 | 9 | 3 | 6 |
| Hymenoptera | Eulophidae | 0 | 133 | 32 | 0 | 12 |
| Hymenoptera | Eupelmidae | 0 | 3 | 9 | 1 | 3 |
| Hymenoptera | Eurytomidae | 0 | 7 | 5 | 0 | 0 |
| Hymenoptera | Evaniidae | 0 | 2 | 13 | 0 | 6 |
| Hymenoptera | Figitidae | 0 | 15 | 5 | 1 | 6 |
| Hymenoptera | Formicidae | 107 | 3743 | 454 | 3547 | 1060 |
| Hymenoptera | Gasteruptiidae | 0 | 0 | 5 | 0 | 0 |
| Hymenoptera | Halictidae | 0 | 1 | 62 | 2 | 14 |
| Hymenoptera | Ichneumonidae | 0 | 123 | 53 | 0 | 11 |
| Hymenoptera | Megachilidae | 0 | 0 | 4 | 1 | 1 |
| Hymenoptera | Mutillidae | 0 | 0 | 7 | 1 | 2 |
| Hymenoptera | Mymaridae | 0 | 13 | 3 | 1 | 5 |
| Hymenoptera | Ormyridae | 0 | 1 | 1 | 0 | 0 |
| Hymenoptera | Pemphredonidae | 0 | 0 | 15 | 0 | 0 |
| Hymenoptera | Philanthidae | 0 | 0 | 2 | 0 | 0 |
| Hymenoptera | Pirenidae | 0 | 1 | 0 | 0 | 0 |
| Hymenoptera | Platygastridae | 0 | 22 | 4 | 1 | 3 |
| Hymenoptera | Pompilidae | 0 | 22 | 69 | 2 | 51 |
| Hymenoptera | Psenidae | 0 | 0 | 8 | 0 | 0 |
| Hymenoptera | Pteromalidae | 0 | 10 | 10 | 2 | 1 |
| Hymenoptera | Scelionidae | 0 | 157 | 80 | 50 | 247 |
| Hymenoptera | Scoliidae | 0 | 1 | 3 | 0 | 1 |
| Hymenoptera | Sphecidae | 0 | 0 | 13 | 1 | 0 |
| Hymenoptera | Tenthredinidae | 0 | 0 | 1 | 1 | 1 |
| Hymenoptera | Tetracampidae | 0 | 0 | 1 | 0 | 0 |
| Hymenoptera | Thynnidae | 0 | 1 | 21 | 0 | 0 |
| Hymenoptera | Tiphiidae | 0 | 0 | 8 | 1 | 11 |
| Hymenoptera | Torymidae | 0 | 4 | 12 | 0 | 1 |
| Hymenoptera | Trichogrammatidae | 0 | 3 | 0 | 0 | 0 |
| Hymenoptera | Vespidae | 0 | 0 | 7 | 1 | 5 |
| Lepidoptera | Adelidae | 0 | 1 | 0 | 0 | 0 |
| Lepidoptera | Argyresthiidae | 0 | 3 | 0 | 0 | 0 |
| Lepidoptera | Autostichidae | 0 | 6 | 0 | 0 | 0 |
| Lepidoptera | Batrachedridae | 0 | 2 | 0 | 0 | 0 |
| Lepidoptera | Bedelliidae | 0 | 2 | 0 | 0 | 0 |
| Lepidoptera | Blastobasidae | 1 | 6 | 4 | 0 | 0 |
| Lepidoptera | Bombycidae | 0 | 1 | 0 | 0 | 0 |
| Lepidoptera | Brachodidae | 0 | 0 | 1 | 0 | 0 |
| Lepidoptera | Bucculatricidae | 0 | 5 | 3 | 0 | 0 |
| Lepidoptera | Choreutidae | 0 | 7 | 0 | 0 | 1 |
| Lepidoptera | Coleophoridae | 0 | 19 | 0 | 0 | 0 |
| Lepidoptera | Cosmopterigidae | 3 | 251 | 43 | 0 | 0 |
| Lepidoptera | Cossidae | 0 | 58 | 0 | 0 | 0 |
| Lepidoptera | Crambidae | 28 | 1066 | 39 | 0 | 0 |
| Lepidoptera | Depressariidae | 3 | 115 | 5 | 0 | 0 |
| Lepidoptera | Dryadaulidae | 0 | 0 | 1 | 0 | 0 |
| Lepidoptera | Dudgeoneidae | 0 | 1 | 0 | 0 | 0 |
| Lepidoptera | Elachistidae | 0 | 4 | 2 | 0 | 0 |
| Lepidoptera | Erebidae | 27 | 3615 | 156 | 1 | 2 |
| Lepidoptera | Eriocottidae | 1 | 318 | 1 | 0 | 0 |
| Lepidoptera | Eupterotidae | 0 | 3 | 0 | 0 | 0 |
| Lepidoptera | Euteliidae | 0 | 23 | 3 | 1 | 0 |
| Lepidoptera | Gelechiidae | 4 | 393 | 34 | 0 | 2 |
| Lepidoptera | Geometridae | 7 | 567 | 15 | 0 | 0 |
| Lepidoptera | Glyphipterigidae | 0 | 5 | 1 | 0 | 0 |
| Lepidoptera | Gracillariidae | 1 | 93 | 26 | 0 | 0 |
| Lepidoptera | Hesperiidae | 0 | 0 | 5 | 0 | 2 |
| Lepidoptera | Immidae | 0 | 26 | 2 | 0 | 0 |
| Lepidoptera | Lasiocampidae | 0 | 79 | 0 | 0 | 0 |
| Lepidoptera | Lecithoceridae | 3 | 131 | 8 | 0 | 1 |
| Lepidoptera | Limacodidae | 0 | 66 | 0 | 0 | 0 |
| Lepidoptera | Lycaenidae | 0 | 3 | 33 | 0 | 3 |
| Lepidoptera | Lyonetiidae | 0 | 0 | 2 | 0 | 0 |
| Lepidoptera | Momphidae | 0 | 2 | 0 | 0 | 0 |
| Lepidoptera | Nepticulidae | 0 | 11 | 9 | 0 | 0 |
| Lepidoptera | Noctuidae | 3 | 1922 | 33 | 4 | 1 |
| Lepidoptera | Nolidae | 0 | 326 | 1 | 0 | 1 |
| Lepidoptera | Notodontidae | 0 | 136 | 0 | 0 | 0 |
| Lepidoptera | Nymphalidae | 0 | 1 | 2 | 1 | 0 |
| Lepidoptera | Oecophoridae | 0 | 4 | 0 | 0 | 0 |
| Lepidoptera | Pieridae | 0 | 6 | 0 | 0 | 11 |
| Lepidoptera | Plutellidae | 0 | 24 | 3 | 0 | 0 |
| Lepidoptera | Praydidae | 0 | 2 | 0 | 0 | 0 |
| Lepidoptera | Psychidae | 1 | 19 | 2 | 0 | 0 |
| Lepidoptera | Pterophoridae | 0 | 16 | 0 | 0 | 0 |
| Lepidoptera | Pyralidae | 8 | 1161 | 11 | 1 | 2 |
| Lepidoptera | Saturniidae | 0 | 51 | 0 | 1 | 0 |
| Lepidoptera | Scythrididae | 1 | 13 | 1 | 0 | 0 |
| Lepidoptera | Sphingidae | 0 | 353 | 0 | 0 | 0 |
| Lepidoptera | Stathmopodidae | 0 | 9 | 0 | 0 | 0 |
| Lepidoptera | Thyrididae | 0 | 20 | 0 | 0 | 0 |
| Lepidoptera | Tineidae | 2 | 58 | 10 | 0 | 0 |
| Lepidoptera | Tortricidae | 15 | 367 | 26 | 0 | 0 |
| Lepidoptera | Uraniidae | 0 | 5 | 3 | 0 | 0 |
| Lepidoptera | Yponomeutidae | 0 | 3 | 0 | 0 | 0 |
| Mantodea | Eremiaphilidae | 0 | 1 | 0 | 0 | 0 |
| Mantodea | Galinthiadidae | 0 | 2 | 0 | 0 | 0 |
| Mantodea | Hoplocoryphidae | 0 | 2 | 0 | 1 | 0 |
| Mantodea | Hymenopodidae | 0 | 22 | 0 | 0 | 0 |
| Mantodea | Mantidae | 0 | 1 | 0 | 0 | 0 |
| Mantodea | Miomantidae | 0 | 1 | 0 | 1 | 0 |
| Mesostigmata | Ameroseiidae | 0 | 2 | 0 | 0 | 0 |
| Mesostigmata | Ascidae | 0 | 1 | 0 | 0 | 0 |
| Mesostigmata | Laelapidae | 0 | 1 | 0 | 0 | 0 |
| Mesostigmata | Macrochelidae | 0 | 3 | 0 | 0 | 0 |
| Mesostigmata | Melicharidae | 0 | 16 | 0 | 0 | 0 |
| Mesostigmata | Pachylaelapidae | 0 | 1 | 0 | 0 | 0 |
| Neuroptera | Berothidae | 0 | 43 | 2 | 0 | 0 |
| Neuroptera | Chrysopidae | 0 | 2 | 1 | 0 | 0 |
| Neuroptera | Coniopterygidae | 0 | 13 | 0 | 0 | 0 |
| Neuroptera | Hemerobiidae | 0 | 2 | 0 | 0 | 0 |
| Neuroptera | Mantispidae | 0 | 13 | 0 | 0 | 0 |
| Neuroptera | Myrmeleontidae | 0 | 2 | 0 | 0 | 1 |
| Neuroptera | Sisyridae | 0 | 1 | 0 | 0 | 0 |
| Odonata | Coenagrionidae | 0 | 1 | 0 | 0 | 0 |
| Odonata | Libellulidae | 0 | 2 | 0 | 0 | 0 |
| Orthoptera | Acrididae | 1 | 92 | 25 | 63 | 30 |
| Orthoptera | Gryllacrididae | 0 | 1 | 0 | 0 | 0 |
| Orthoptera | Gryllidae | 2 | 214 | 2 | 196 | 22 |
| Orthoptera | Gryllotalpidae | 0 | 68 | 0 | 0 | 0 |
| Orthoptera | Oecanthidae | 0 | 12 | 0 | 0 | 1 |
| Orthoptera | Pyrgomorphidae | 0 | 1 | 0 | 7 | 2 |
| Orthoptera | Tetrigidae | 0 | 167 | 3 | 10 | 11 |
| Orthoptera | Tettigoniidae | 0 | 125 | 6 | 3 | 7 |
| Orthoptera | Tridactylidae | 0 | 132 | 0 | 3 | 6 |
| Orthoptera | Trigonidiidae | 4 | 377 | 0 | 15 | 10 |
| Plecoptera | Perlidae | 0 | 1 | 0 | 0 | 0 |
| Poduromorpha | Hypogastruridae | 0 | 0 | 0 | 1 | 0 |
| Pseudoscorpiones | Olpiidae | 0 | 2 | 0 | 0 | 0 |
| Psocodea | Amphipsocidae | 0 | 1 | 0 | 0 | 0 |
| Psocodea | Caeciliusidae | 1 | 4 | 8 | 0 | 0 |
| Psocodea | Ectopsocidae | 0 | 1 | 33 | 1 | 0 |
| Psocodea | Lepidopsocidae | 0 | 0 | 21 | 1 | 0 |
| Psocodea | Liposcelididae | 0 | 1 | 0 | 0 | 0 |
| Psocodea | Peripsocidae | 0 | 1 | 0 | 0 | 0 |
| Psocodea | Philotarsidae | 0 | 2 | 1 | 0 | 0 |
| Psocodea | Prionoglarididae | 0 | 1 | 0 | 0 | 0 |
| Psocodea | Pseudocaeciliidae | 0 | 0 | 8 | 0 | 0 |
| Strepsiptera | Corioxenidae | 0 | 1 | 0 | 0 | 0 |
| Strepsiptera | Mengenillidae | 0 | 16 | 0 | 0 | 0 |
| Strepsiptera | Myrmecolacidae | 0 | 1 | 0 | 0 | 0 |
| Thysanoptera | Aeolothripidae | 0 | 69 | 0 | 0 | 0 |
| Thysanoptera | Phlaeothripidae | 0 | 6 | 4 | 2 | 5 |

Supplementary Table 3: The number of crop pest species detected in each trap type.

| **Order** | **Family** | **Species** | **CDC** | **Heath** | **Malaise** | **Pitfall** | **Yellow Pan** |
| --- | --- | --- | --- | --- | --- | --- | --- |
| Coleoptera | Bostrichidae | *Apate monachus* | 0 | 0 | 1 | 0 | 0 |
| Coleoptera | Brentidae | *Cylas brunneus* | 0 | 1 | 0 | 0 | 0 |
| Coleoptera | Brentidae | *Cylas puncticollis* | 0 | 2 | 0 | 0 | 0 |
| Coleoptera | Chrysomelidae | *Monolepta jacksoni* | 0 | 74 | 0 | 0 | 0 |
| Coleoptera | Curculionidae | *Euplatypus hintzi* | 0 | 208 | 0 | 0 | 0 |
| Coleoptera | Meloidae | *Hycleus hermanniae* | 1 | 0 | 0 | 50 | 23 |
| Coleoptera | Tenebrionidae | *Gonocephalum simplex* | 0 | 0 | 0 | 1 | 0 |
| Diptera | Drosophilidae | *Zaprionus indianus* | 0 | 0 | 4 | 0 | 0 |
| Hemiptera | Aleyrodidae | *Aleurodicus dispersus* | 0 | 9 | 0 | 0 | 0 |
| Hemiptera | Aphididae | *Aphis nerii* | 0 | 3 | 0 | 1 | 4 |
| Hemiptera | Aphididae | *Aphis spiraecola* | 0 | 0 | 0 | 0 | 1 |
| Hemiptera | Aphididae | *Rhopalosiphum rufiabdominale* | 0 | 11 | 0 | 0 | 1 |
| Hemiptera | Aphrophoridae | *Poophilus costalis* | 0 | 1 | 0 | 0 | 0 |
| Hemiptera | Delphacidae | *Peregrinus maidis* | 0 | 106 | 2 | 1 | 7 |
| Hemiptera | Miridae | *Nesidiocoris tenuis* | 0 | 19 | 0 | 0 | 0 |
| Hemiptera | Pentatomidae | *Nezara viridula* | 0 | 1 | 0 | 0 | 0 |
| Lepidoptera | Crambidae | *Duponchelia fovealis* | 0 | 1 | 1 | 0 | 0 |
| Lepidoptera | Crambidae | *Herpetogramma licarsisalis* | 0 | 3 | 0 | 0 | 0 |
| Lepidoptera | Crambidae | *Leucinodes africensis* | 0 | 1 | 0 | 0 | 0 |
| Lepidoptera | Gelechiidae | *Tuta absoluta* | 0 | 3 | 0 | 0 | 0 |
| Lepidoptera | Noctuidae | *Spodoptera frugiperda* | 0 | 2 | 0 | 0 | 0 |
| Lepidoptera | Noctuidae | *Spodoptera littoralis* | 0 | 2 | 0 | 4 | 0 |
| Lepidoptera | Pyralidae | *Etiella zinckenella* | 0 | 4 | 0 | 0 | 0 |
| Lepidoptera | Tortricidae | *Thaumatotibia leucotreta* | 0 | 25 | 1 | 0 | 0 |
| Thysanoptera | Thripidae | *Ayyaria chaetophora* | 0 | 0 | 0 | 0 | 4 |

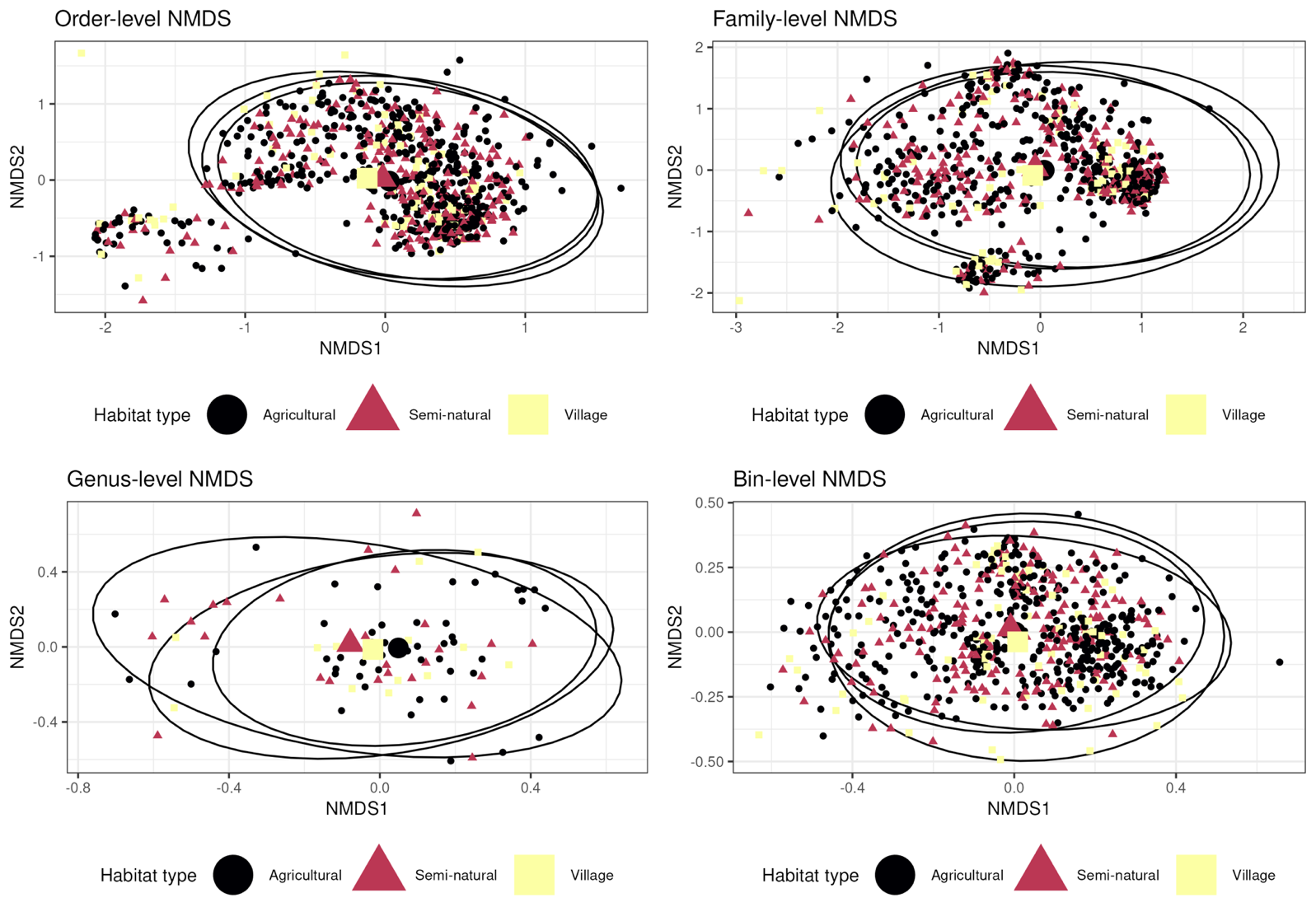

Supplementary figure 4: NMDS plots comparing insect assemblages among different habitat types. Each plot differs in terms of the taxonomic resolution at which samples were categorised. The larger points are the centroids for each trap type.

Supplementary Table 4: The number of individuals of Dipteran families known to contain blood-feeders detected in each trap type.

| **Order** | **Family** | **CDC** | **Heath** | **Malaise** | **Pitfall** | **Yellow Pan** |
| --- | --- | --- | --- | --- | --- | --- |
| Diptera | Ceratopogonidae | 57 | 1037 | 442 | 1 | 19 |
| Diptera | Culicidae | 180 | 6 | 4 | 1 | 1 |
| Diptera | Simuliidae | 0 | 10 | 2 | 0 | 0 |
| Diptera | Tabanidae | 3 | 36 | 21 | 0 | 0 |

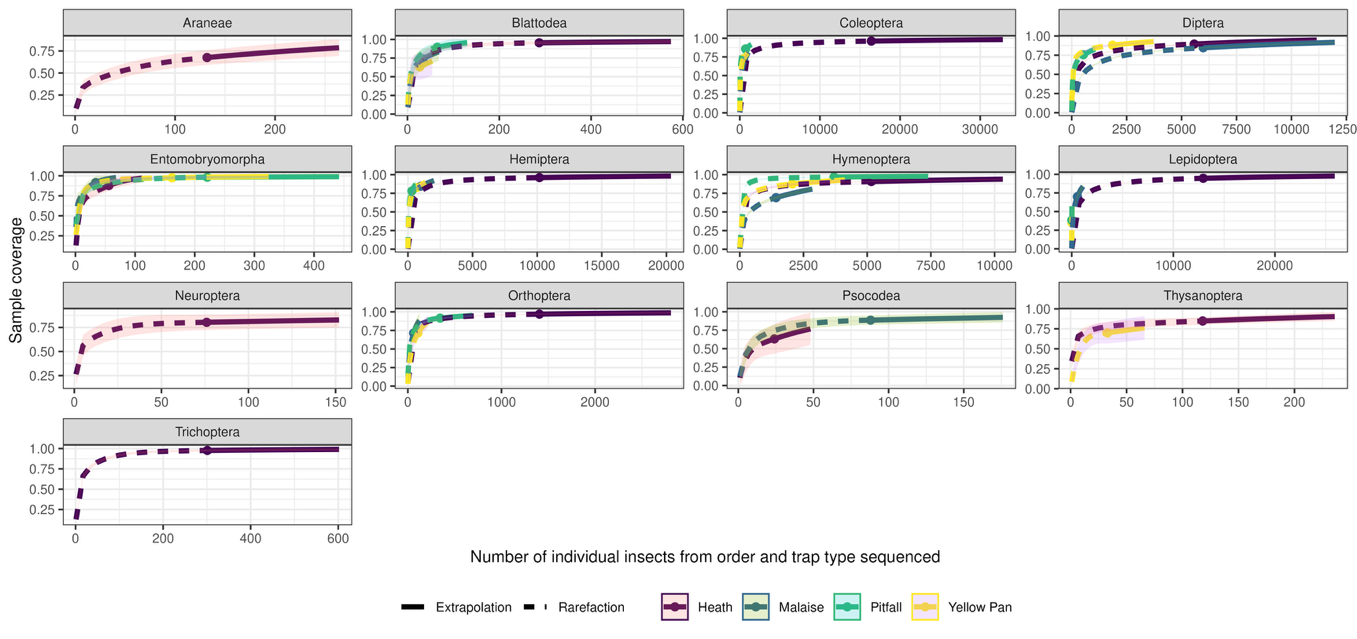

Supplementary Figure 5: type 2 iNEXT plot, showing the estimated sample coverage for each taxonomic order and trap type.

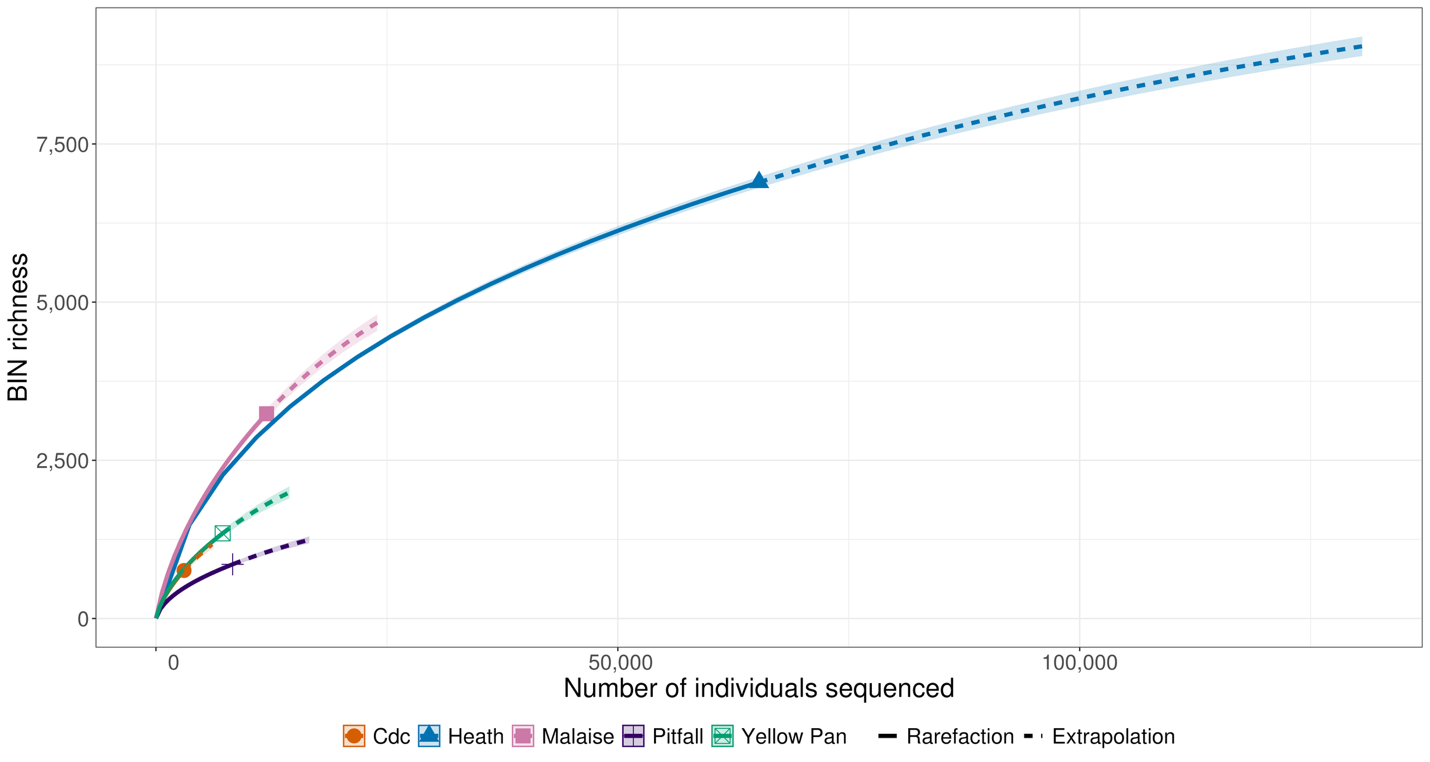

Supplementary Figure 6: Type 1 iNEXT plot showing the observed and extrapolated accumulation of BINs in relation to the number of individuals sequenced for each trap type.

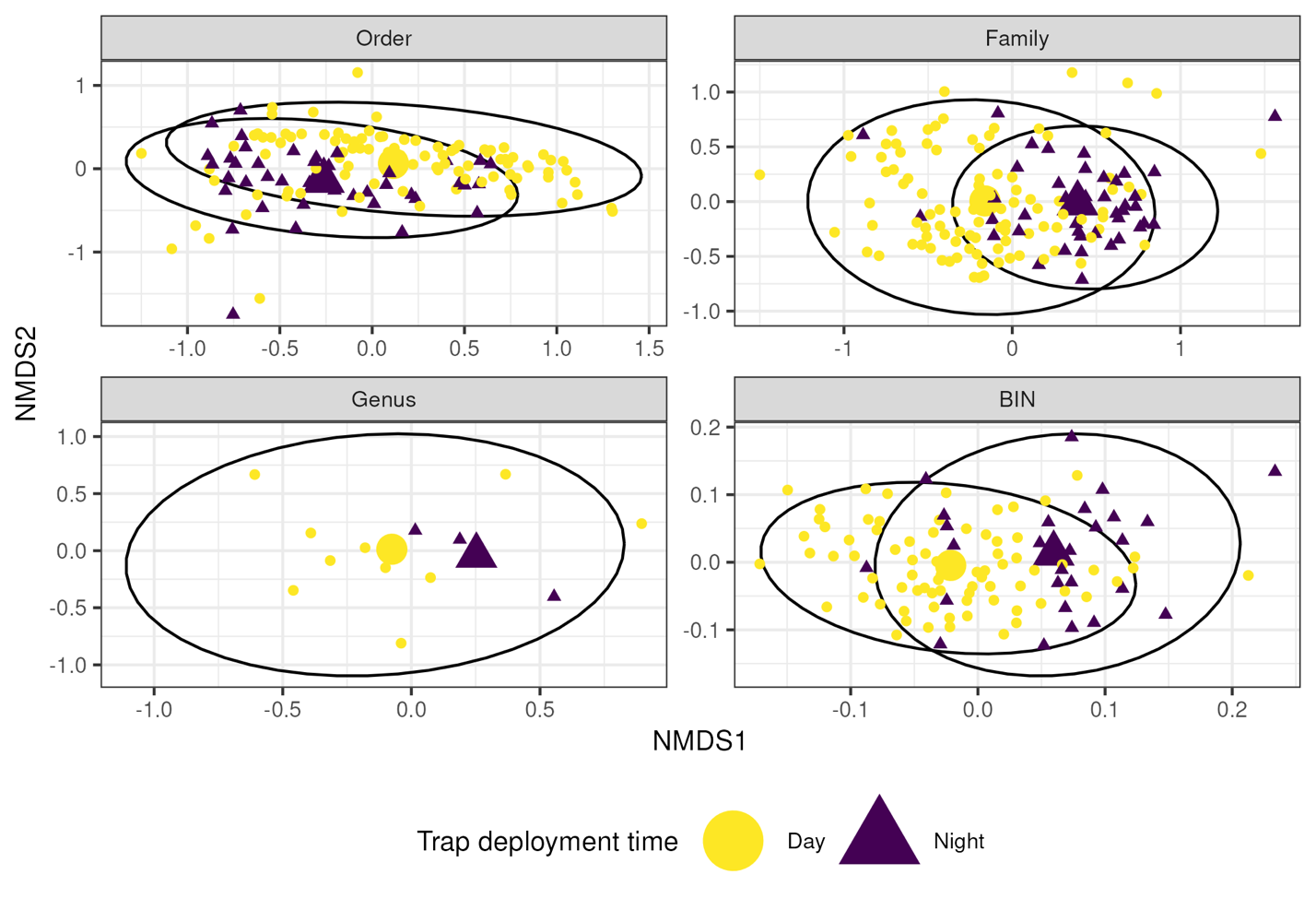

Supplementary Figure 7: NMDS plots comparing insect assemblages among day and night-deployed malaise traps. Each plot differs in terms of the taxonomic resolution at which samples were categorised. The larger points are the centroids for each trap type. There are fewer data points for the genus-level plot because many samples were not assigned to genus level, despite being identified to family level and assigned a BIN.

Supplementary Table 5: the most abundant 20 taxonomic families for each trap type, and whether they were found in Srivathsan et al.’s 2023 paper documenting the globally most abundant taxonomic families in Malaise trap data. E.g. Cecidomyiidae were the most abundant family in CDC data, and were found in Srivathsan et al, Drosophilidae were 11^th^ most abundant family in Malaise trap data and were not found in Srivathsan et al.’s 20 most abundant families in Malaise trap data.

|  |  | **Trap abundance ranking** | | | | |  |
| --- | --- | --- | --- | --- | --- | --- | --- |
| **Order** | **Family** | **CDC** | **Yellow Pan** | **Pitfall** | **Malaise** | **Heath** | **In Srivathsan et al** |
| Diptera | Cecidomyiidae | 1 | 14 | 17 | 1 | 16 | TRUE |
| Diptera | Culicidae | 2 |  |  |  |  | FALSE |
| Hymenoptera | Formicidae | 3 | 1.5 | 1 | 5 | 3 | TRUE |
| Diptera | Ceratopogonidae | 4 |  |  | 6 | 15 | TRUE |
| Coleoptera | Phalacridae | 5 |  |  |  |  | FALSE |
| Hemiptera | Cicadellidae | 6 | 3 | 4 | 3 | 4 | TRUE |
| Diptera | Limoniidae | 7 |  |  |  |  | FALSE |
| Diptera | Muscidae | 8 |  | 20 | 4 |  | TRUE |
| Lepidoptera | Crambidae | 9.5 |  |  |  | 14 | TRUE |
| Diptera | Drosophilidae | 9.5 |  |  | 11 |  | FALSE |
| Lepidoptera | Erebidae | 11 |  |  | 15 | 2 | TRUE |
| Diptera | Psychodidae | 12.5 |  |  | 13 |  | TRUE |
| Coleoptera | Staphylinidae | 12.5 | 18 | 8 |  | 1 | FALSE |
| Diptera | Sciaridae | 14 |  |  | 10 | 17 | TRUE |
| Coleoptera | Chrysomelidae | 15 | 7 | 10 | 19 | 13 | FALSE |
| Diptera | Calliphoridae | 16 |  |  | 16 |  | FALSE |
| Coleoptera | Scarabaeidae | 17 |  |  |  | 9 | FALSE |
| Coleoptera | Curculionidae | 18.5 |  |  |  |  | FALSE |
| Lepidoptera | Tortricidae | 18.5 |  |  |  |  | FALSE |
| Coleoptera | Carabidae | 20 |  | 6 |  | 5 | FALSE |
| Diptera | Dolichopodidae |  | 1.5 | 12 | 18 |  | TRUE |
| Hymenoptera | Crabronidae |  | 4 |  |  |  | FALSE |
| Hymenoptera | Scelionidae |  | 5 | 18 |  |  | FALSE |
| Diptera | Sarcophagidae |  | 6 |  | 12 |  | FALSE |
| Hymenoptera | Bethylidae |  | 8 |  |  |  | TRUE |
| Diptera | Chloropidae |  | 9 | 19 | 2 |  | TRUE |
| Hemiptera | Aphididae |  | 10 |  |  |  | FALSE |
| Diptera | Phoridae |  | 11 | 7 | 8 |  | TRUE |
| Entomobryomorpha | Isotomidae |  | 12 | 5 |  |  | FALSE |
| Hemiptera | Delphacidae |  | 13 |  |  | 8 | FALSE |
| Hymenoptera | Diapriidae |  | 15 |  |  |  | FALSE |
| Coleoptera | Meloidae |  | 16 | 3 |  |  | FALSE |
| Hymenoptera | Pompilidae |  | 17 |  |  |  | FALSE |
| Diptera | Milichiidae |  | 19 |  | 7 |  | FALSE |
| Orthoptera | Gryllidae |  |  | 2 |  |  | FALSE |
| Diptera | Sphaeroceridae |  |  | 9 | 20 |  | TRUE |
| Coleoptera | Nitidulidae |  |  | 11 |  |  | FALSE |
| Orthoptera | Acrididae |  |  | 13 |  |  | FALSE |
| Coleoptera | Anthicidae |  |  | 14 |  |  | FALSE |
| Blattodea | Termitidae |  |  | 15 |  |  | FALSE |
| Diptera | Ephydridae |  |  | 16 |  |  | FALSE |
| Diptera | Lonchaeidae |  |  |  | 9 |  | FALSE |
| Hymenoptera | Braconidae |  |  |  | 14 | 20 | TRUE |
| Diptera | Chironomidae |  |  |  | 17 | 18 | TRUE |
| Lepidoptera | Noctuidae |  |  |  |  | 6 | FALSE |
| Coleoptera | Hydrophilidae |  |  |  |  | 7 | FALSE |
| Coleoptera | Dytiscidae |  |  |  |  | 10 | FALSE |
| Lepidoptera | Pyralidae |  |  |  |  | 11 | FALSE |
| Hemiptera | Rhyparochromidae |  |  |  |  | 12 | FALSE |
| Lepidoptera | Geometridae |  |  |  |  | 19 | FALSE |

Supplementary Table 6: the numbers of individuals of different arthropod taxa collected in Malaise traps during daytime and nighttime.

| Order | Family | Day | Night |
| --- | --- | --- | --- |
| Araneae | Araneidae | 0 | 0 |
| Araneae | Cheiracanthiidae | 0 | 0 |
| Araneae | Lycosidae | 0 | 0 |
| Araneae | Theridiidae | 0 | 0 |
| Araneae | Thomisidae | 0 | 0 |
| Blattodea | Blattellidae | 0 | 0 |
| Blattodea | Ectobiidae | 0 | 1 |
| Blattodea | Kalotermitidae | 0 | 0 |
| Blattodea | Rhinotermitidae | 0 | 0 |
| Blattodea | Termitidae | 5 | 0 |
| Coleoptera | Aderidae | 2 | 2 |
| Coleoptera | Anthicidae | 6 | 3 |
| Coleoptera | Anthribidae | 7 | 3 |
| Coleoptera | Biphyllidae | 0 | 1 |
| Coleoptera | Bostrichidae | 2 | 1 |
| Coleoptera | Brentidae | 0 | 0 |
| Coleoptera | Buprestidae | 4 | 0 |
| Coleoptera | Carabidae | 5 | 3 |
| Coleoptera | Cerambycidae | 4 | 2 |
| Coleoptera | Cerylonidae | 0 | 0 |
| Coleoptera | Chrysomelidae | 48 | 21 |
| Coleoptera | Cleridae | 1 | 1 |
| Coleoptera | Coccinellidae | 13 | 6 |
| Coleoptera | Corylophidae | 3 | 0 |
| Coleoptera | Cryptophagidae | 0 | 0 |
| Coleoptera | Curculionidae | 9 | 1 |
| Coleoptera | Dermestidae | 0 | 0 |
| Coleoptera | Dytiscidae | 0 | 0 |
| Coleoptera | Elateridae | 2 | 0 |
| Coleoptera | Elmidae | 0 | 0 |
| Coleoptera | Erotylidae | 1 | 0 |
| Coleoptera | Euxestidae | 0 | 0 |
| Coleoptera | Heteroceridae | 0 | 0 |
| Coleoptera | Histeridae | 0 | 0 |
| Coleoptera | Hybosoridae | 0 | 0 |
| Coleoptera | Hydraenidae | 0 | 0 |
| Coleoptera | Hydrophilidae | 0 | 0 |
| Coleoptera | Laemophloeidae | 3 | 0 |
| Coleoptera | Lampyridae | 0 | 0 |
| Coleoptera | Latridiidae | 1 | 0 |
| Coleoptera | Leiodidae | 0 | 0 |
| Coleoptera | Limnichidae | 0 | 0 |
| Coleoptera | Lycidae | 1 | 1 |
| Coleoptera | Meloidae | 3 | 0 |
| Coleoptera | Melyridae | 3 | 0 |
| Coleoptera | Mordellidae | 9 | 3 |
| Coleoptera | Mycetophagidae | 0 | 0 |
| Coleoptera | Nitidulidae | 8 | 0 |
| Coleoptera | Noteridae | 0 | 0 |
| Coleoptera | Phalacridae | 4 | 6 |
| Coleoptera | Ptinidae | 1 | 0 |
| Coleoptera | Scarabaeidae | 1 | 1 |
| Coleoptera | Scirtidae | 0 | 0 |
| Coleoptera | Scraptiidae | 3 | 1 |
| Coleoptera | Silvanidae | 0 | 0 |
| Coleoptera | Spercheidae | 0 | 0 |
| Coleoptera | Staphylinidae | 7 | 2 |
| Coleoptera | Tenebrionidae | 3 | 0 |
| Coleoptera | Throscidae | 2 | 0 |
| Coleoptera | Zopheridae | 0 | 0 |
| Dermaptera | Spongiphoridae | 1 | 0 |
| Diptera | Agromyzidae | 50 | 4 |
| Diptera | Anthomyiidae | 3 | 1 |
| Diptera | Bombyliidae | 3 | 2 |
| Diptera | Calliphoridae | 51 | 9 |
| Diptera | Cecidomyiidae | 458 | 226 |
| Diptera | Ceratopogonidae | 281 | 68 |
| Diptera | Chaoboridae | 0 | 0 |
| Diptera | Chironomidae | 53 | 42 |
| Diptera | Chloropidae | 496 | 62 |
| Diptera | Chyromyidae | 3 | 1 |
| Diptera | Culicidae | 1 | 3 |
| Diptera | Diopsidae | 1 | 0 |
| Diptera | Dolichopodidae | 86 | 8 |
| Diptera | Drosophilidae | 103 | 22 |
| Diptera | Ephydridae | 28 | 4 |
| Diptera | Hybotidae | 59 | 5 |
| Diptera | Keroplatidae | 2 | 0 |
| Diptera | Lauxaniidae | 25 | 8 |
| Diptera | Limoniidae | 4 | 4 |
| Diptera | Lonchaeidae | 119 | 4 |
| Diptera | Milichiidae | 175 | 19 |
| Diptera | Muscidae | 346 | 50 |
| Diptera | Mycetophilidae | 12 | 5 |
| Diptera | Periscelididae | 2 | 1 |
| Diptera | Phoridae | 165 | 18 |
| Diptera | Pipunculidae | 11 | 1 |
| Diptera | Platystomatidae | 52 | 1 |
| Diptera | Psychodidae | 94 | 26 |
| Diptera | Pyrgotidae | 0 | 1 |
| Diptera | Sarcophagidae | 59 | 17 |
| Diptera | Sciaridae | 134 | 38 |
| Diptera | Simuliidae | 2 | 0 |
| Diptera | Sphaeroceridae | 70 | 7 |
| Diptera | Stratiomyidae | 29 | 3 |
| Diptera | Syrphidae | 18 | 3 |
| Diptera | Tabanidae | 10 | 0 |
| Diptera | Tachinidae | 78 | 6 |
| Diptera | Tephritidae | 7 | 0 |
| Diptera | Ulidiidae | 26 | 6 |
| Entomobryomorpha | Entomobryidae | 14 | 7 |
| Entomobryomorpha | Isotomidae | 0 | 0 |
| Entomobryomorpha | Seiridae | 0 | 0 |
| Ephemeroptera | Baetidae | 0 | 0 |
| Hemiptera | Achilidae | 0 | 0 |
| Hemiptera | Aleyrodidae | 14 | 2 |
| Hemiptera | Alydidae | 0 | 0 |
| Hemiptera | Anthocoridae | 0 | 0 |
| Hemiptera | Aphididae | 19 | 11 |
| Hemiptera | Aphrophoridae | 0 | 2 |
| Hemiptera | Aradidae | 0 | 0 |
| Hemiptera | Belostomatidae | 0 | 0 |
| Hemiptera | Ceratocombidae | 0 | 1 |
| Hemiptera | Cicadellidae | 327 | 234 |
| Hemiptera | Cixiidae | 0 | 0 |
| Hemiptera | Coreidae | 0 | 0 |
| Hemiptera | Corixidae | 0 | 0 |
| Hemiptera | Cydnidae | 0 | 0 |
| Hemiptera | Cymidae | 0 | 0 |
| Hemiptera | Delphacidae | 6 | 2 |
| Hemiptera | Derbidae | 0 | 1 |
| Hemiptera | Dictyopharidae | 0 | 0 |
| Hemiptera | Flatidae | 1 | 1 |
| Hemiptera | Fulgoridae | 0 | 0 |
| Hemiptera | Gerridae | 0 | 0 |
| Hemiptera | Hebridae | 0 | 0 |
| Hemiptera | Hydrometridae | 0 | 0 |
| Hemiptera | Issidae | 0 | 1 |
| Hemiptera | Largidae | 1 | 0 |
| Hemiptera | Lethaeidae | 0 | 0 |
| Hemiptera | Lygaeidae | 6 | 2 |
| Hemiptera | Meenoplidae | 5 | 5 |
| Hemiptera | Mesoveliidae | 0 | 0 |
| Hemiptera | Miridae | 15 | 4 |
| Hemiptera | Nabidae | 0 | 0 |
| Hemiptera | Notonectidae | 0 | 0 |
| Hemiptera | Ochteridae | 0 | 0 |
| Hemiptera | Pentatomidae | 0 | 0 |
| Hemiptera | Pleidae | 0 | 0 |
| Hemiptera | Psyllidae | 12 | 4 |
| Hemiptera | Psylloidea_incertae_sedis | 2 | 0 |
| Hemiptera | Pyrrhocoridae | 0 | 0 |
| Hemiptera | Reduviidae | 2 | 0 |
| Hemiptera | Rhyparochromidae | 2 | 3 |
